# Cortisol Drives Pregnancy-Associated Induction of Hepatic OAT2, NTCP, and OCT1 in HepaRG cells Through GR-, HNF1α-, and HNF4α-Dependent Signaling

**DOI:** 10.64898/2026.06.15.732466

**Authors:** Sejal Sharma, Yik Pui Tsang, Jashvant D. Unadkat

## Abstract

Pregnancy induces or represses hepatic drug metabolism. Whether pregnancy affects hepatic drug transport is unexplored. We previously showed that a cocktail of pregnancy-related hormones (PRHC) induces mRNA expression and activity of sodium/taurocholate cotransporting polypeptide (NTCP), organic anion transporter 2 (OAT2), and organic cation transporter 1 (OCT1, mRNA only) in differentiated HepaRG cells. Here, using HepaRG cells, we identified cortisol as the hormone primarily responsible for this induction and explored the underlying mechanisms. Clustered regularly interspaced short palindromic repeats (CRISPR)-Cas9-mediated knockdown studies in HepaRG cells showed that the glucocorticoid receptor (GR) is the primary mediator of this response. GR knockdown markedly attenuated cortisol-induced NTCP, OAT2, and OCT1 mRNA expression and activity. Cortisol also induced the mRNA expression of regulatory factors, including pregnane X receptor (PXR), constitutive androstane receptor (CAR), and hepatocyte nuclear factor (HNF) 4 alpha (HNF4α). HNF4α knockdown selectively attenuated OAT2 and OCT1 induction, whereas HNF1α knockdown enhanced NTCP induction, attenuated OCT1 induction, and reduced basal organic anion transporting polypeptide 1B1 (OATP1B1) expression. In contrast, knockdown of CAR or PXR did not significantly alter cortisol-mediated transporter regulation. These data identify cortisol as the principal PRH driving regulation of the hepatic OAT2, NTCP, and OCT1 in HepaRG cells and indicate that this response is mediated primarily by GR, with selective downstream contributions from HNF4α and HNF1α. These findings provide mechanistic insights into pregnancy-associated changes in hepatic transporter-mediated drug disposition, including when antenatal corticosteroids are administered to pregnant women to prevent respiratory distress syndrome in their prematurely born infants.

**Significance Statement:** The extent and mechanisms by which pregnancy-related hormones regulate hepatic uptake transporters remain poorly defined. This study identifies cortisol as the principal pregnancy-related hormone driving NTCP, OAT2, and OCT1 induction in HepaRG cells and shows that this response is mediated primarily through GR, with transporter-specific contributions from HNF4α and HNF1α.

## 1. Introduction

Pregnancy produces coordinated physiological and hormonal changes that markedly alter drug disposition ^1^. In the liver, activity of drug metabolizing enzymes, most notably cytochrome P450 (CYP) 3A, increases during gestation and has been linked to subtherapeutic concentrations of anti-retroviral drug indinavir ^2,3^. Emerging evidence indicates that hepatic drug transporters are also modulated during pregnancy ^4,5^, yet perpetrator hormone(s) and regulatory mechanism(s) remain undefined.

The pregnancy-related hormonal milieu is characterized by gestational-age dependent increases in plasma concentrations of estrogens (estrone [E1], estradiol [E2], estriol [E3], and estetrol [E4]), progesterone, cortisol, oxytocin, and placental growth hormone (PGH) ^5,6^. In human hepatocytes from premenopausal female donors, a pregnancy-related hormone cocktail (PRHC) formulated to match average third-trimester (T3) plasma concentrations of these eight hormones increased the expression and/or activity of several hepatic transporters, with the magnitude of induction following the rank order: sodium/taurocholate cotransporting polypeptide (NTCP) ≈ organic anion transporter 2 (OAT2) ≈ organic cation transporter 1 (OCT1) > organic anion transporting polypeptide 2B1 (OATP2B1), whereas OATP1B3 was repressed ^5^. We observed a similar pattern of effect of 1x PRHC on the mRNA expression of the uptake transporters in HepaRG cells, except that mRNA expression of OATP1B1 (not OATP1B3) was repressed and that of organic anion transporter 7 (OAT7, not measured in hepatocytes) was induced ^7^.

Moreover, transporter induction by PRHC was positively associated with changes in regulatory factors such as pregnane X receptor (PXR), constitutive androstane receptor (CAR), and hepatocyte nuclear factor (HNF) 4α ^7^. Together, these findings suggest that PRHs modulate the expression and activity of these hepatic transporters, potentially through coordinated changes in regulatory factors. The similar pattern observed in HepaRG cells and human hepatocytes supports the use of HepaRG cells as a mechanistic model to define the hormone-specific transcriptional pathways underlying pregnancy-associated hepatic transporter regulation.

This study aimed to identify the hormone(s), within the PRHC, primarily responsible for induction (or repression) of the above referenced hepatic uptake transporters in HepaRG cells ^7^ and to delineate the regulatory pathways involved. We first compared the effect of PRHC with its individual constituent hormones at physiologic (1×) and supraphysiologic (10×) T3 concentrations on changes in transporter (mRNA and activity) and regulatory factor mRNA expression in HepaRG cells. We then used clustered regularly interspaced short palindromic repeats (CRISPR)/Cas9-mediated knockdown of glucocorticoid receptor (GR), CAR, PXR, HNF4α, and HNF1α to determine their contributions to cortisol-mediated changes in transporter mRNA expression and, for selected transporters, functional activity. This experimental strategy tested the hypothesis that pregnancy-like hepatic transporter induction is driven predominantly by cortisol and is mediated through GR-dependent signaling involving one or more downstream hepatic regulatory factors.

## 2. Materials and Methods

### 2.1 Chemicals and reagents

Differentiated HepaRG cells (passage 16), and HepaRG Thawing and Plating Medium Supplement with Antibiotics were purchased from Biopredic Inc (Nashville, TN, USA). Estrone (E1), estradiol (E2), estriol (E3), estetrol (E4), hydrocortisone (cortisol), progesterone, testosterone, oxytocin, and dimethyl sulfoxide (DMSO) were purchased from MilliporeSigma (Burlington, MA, USA). Tris-HCl, sodium dodecyl sulfate (SDS), calcium chloride (anhydrous), magnesium chloride hexahydrate (MgClß·6HßO), magnesium sulfate heptahydrate (MgSOß·7HßO), potassium chloride (KCl), monopotassium phosphate (KHßPOß), potassium bicarbonate (KHCOß), choline chloride, dipotassium phosphate (KßHPOß), D-glucose, PureLink RNA Mini Kit, High-Capacity cDNA Reverse Transcription Kit, GlutaMAX Supplement (100X), William’s E Medium (no phenol red), Dulbecco’s phosphate-buffered saline with calcium and magnesium (DPBS++), Hanks’ balanced salt solution with calcium and magnesium (HBSS++), Insulin-Transferrin-Selenium (ITS-G; 100X), TrueGuide Synthetic single guide RNAs (sgRNAs; targeting *NR3C1*, *HNF1A*, *HNF4A*, *NR1I2*, and *NR1I3*) (sgRNA sequences shown in Supplementary Table 1), Lipofectamine CRISPRMAX Cas9 Transfection Reagent, TrueCut Cas9 Protein v2, Tris-EDTA buffer, Opti-MEM I Reduced Serum Medium, penicillin-streptomycin (10,000 U/ml; 100X), Pierce Bicinchoninic Acid (BCA) Protein Assay Kit, PureLink PCR Purification Kit, AmpliTaq Gold 360 Master Mix, PCR primers (Supplementary Table 2), TaqMan Fast Advanced Master Mix, TaqMan gene expression assays, GeneArt Genomic Cleavage Detection Kit, DNase I, agarose, UltraPure TBE Buffer (10×), and SYBR Gold Nucleic Acid Gel Stain (10,000X Concentrate in DMSO) were purchased from Thermo Fisher Scientific (Waltham, MA, USA). Genomic DNA Clean & Concentrator-10 was purchased from Zymo Research (Irvine, CA, USA). Placental growth hormone (PGH) was purchased from R&D Systems (Minneapolis, MN, USA). BioCoat Collagen I 24-well and 96-well tissue culture-treated plates were purchased from Corning (Corning, NY, USA). Ecoscint ORIGINAL was purchased from National Diagnostics (Atlanta, GA, USA). [^3^H] taurocholic acid (TA, 50 Ci/mmol, 20 μM) and [^3^H] cGMP (25 Ci/mmol, 20 μM) were purchased from American Radiolabeled Chemicals (St. Louis, MO, USA). [^14^C] metformin (0.114 Ci/mmol, 8.8 mM) was purchased from Moravek (Brea, CA, USA). Sulfobromophthalein (BSP) disodium salt and pyrimethamine were purchased from MedChemExpress (Monmouth Junction, NJ, USA).

The TaqMan Gene Expression Assays used were: *SLC10A1* (NTCP; Hs00161820_m1), *SLC22A7* (OAT2; Hs00198527_m1), *SLC22A1* (OCT1; Hs00427552_m1), *SLC22A9* (OAT7; Hs00971064_m1), *SLCO2B1* (OATP2B1; Hs01030343_m1), *SLCO1B1* (OATP1B1; Hs00272374_m1), *NR1I3* (CAR; Hs00901571_m1), *NR1I2* (PXR; Hs01114267_m1), *HNF4A* (HNF4α; Hs00230853_m1), *HNF1A* (HNF1α; Hs00167041_m1), *NR1H4* (Farnesoid X receptor, FXR; Hs00231968_m1), *AHR* (Aryl Hydrocarbon Receptor, AhR; Hs00169233_m1), *NR3C1* (GR; Hs00353740_m1), *CEBPB* (CCAAT/enhancer-binding protein beta, C/EBPβ; Hs00942496_s1), *NR0B2* (Small Heterodimer Partner, SHP; Hs00222677_m1), *PPARGC1A* (Peroxisome proliferator-activated receptor gamma coactivator 1-alpha, PGC-1α; Hs00173304_m1), *NR5A2* (Liver receptor homolog-1, LRH-1; Hs00187067_m1), *FOXA2* (Forkhead box protein A2, HNF3β; Hs05036278_s1), *CYP3A4* (CYP3A4; Hs00604506_m1), *TAT* (TAT; Hs00356930_m1), *UGT1A1* (UGT1A1; Hs02511055_s1), *CYP2B6* (CYP2B6; Hs03044636_m1) and *GAPDH* (GAPDH; Hs02786624_g1).

### 2.2 HepaRG cell culture and plating

Differentiated HepaRG cells (passage 16) were plated on collagen I-coated plates. For CRISPR transfection and qPCR studies, cells were seeded in 24-well plates at 210,000 cells/cm² and allowed to attach for 24 h in vendor plating medium supplemented with antibiotics. For transporter activity studies, cells were plated on collagen I-coated 96-well plates at 210,000 cells/cm^2^. Media were then replaced with dexamethasone-free maintenance medium consisting of William’s Medium E supplemented with 1× GlutaMAX, 1× ITS-G, and 1× penicillin-streptomycin for 72 h before hormone treatment. Unless otherwise stated, cells were maintained at 37°C in 5% CO_2_, and media were refreshed every 24 h.

### 2.3 Pregnancy-related hormone treatment to identify the hormone driving the changes in transporter expression in HepaRG cells

After the 72-h maintenance period, HepaRG cells were treated with vehicle (0.1% v/v DMSO), a pregnancy-related hormone cocktail (PRHC), or individual pregnancy-related hormones. The PRHC comprised estrone (E1), estradiol (E2), estriol (E3), estetrol (E4), cortisol, progesterone, testosterone, oxytocin, and placental growth hormone (PGH). For the PRHC and individual-hormone experiments, target concentrations were based on the geometric-mean third-trimester (T3) plasma concentrations previously reported in the literature ^5^. To account for hormone depletion in culture, the concentrations added to the medium were adjusted such that the estimated time-averaged hormone concentration over each treatment interval approximated the target 1× or 10× T3 plasma concentration (Table 1). Cells were treated using an 8-h/16-h refreshment schedule as previously described, and this schedule was repeated during the 72-h treatment period ^5,7^. Vehicle-treated cells were refreshed on the same schedule. In experiments where indicated, pooled estrogens (E1+E2+E3+E4) were also tested. The concentrations used for each hormone are provided in Table 1. For all cortisol-only experiments, including the cortisol time-course and the subsequent cortisol-treated CRISPR-Cas9 experiments, medium was refreshed every 24 h because cortisol was not depleted in culture (Table 1).

**Table 1.** Observed steady-state total plasma concentrations of PRHs in the third trimester of pregnancy (T3) and the 1× PRH concentrations used in this study after accounting for hormone depletion in culture. Table adapted from Sharma *et al*., 2025.

| Pregnancy-related hormones (PRH) | T3 total plasma concentration (ng/ml) <sup>a</sup> | 1x PRH concentrations in the media after accounting for depletion (ng/ml) <sup>b</sup> |  |
| --- | --- | --- | --- |
|  |  | 8 hours | 16 hours |
| Estrone (E1) | 9.75 | 30.50 | 61.0 |
| Estradiol (E2) | 13.97 | 44.00 | 88.00 |
| Estriol (E3) | 8.68 | 36.00 | 72.00 |
| Estetrol (E4) | 0.78 | 2.15 | 4.30 |
| Progesterone | 140.11 | 550 | 1,100 |
| Cortisol | 312.36 | 320.00 |  |
| Testosterone | 1.30 | 5.00 | 10.00 |
| Oxytocin | 0.23 | 0.23 |  |
| Placental growth hormone (PGH) | 12.78 | 12.80 |  |

<sup>a</sup>Data are geometric means of total plasma concentration of pregnancy-related hormones <sup>5</sup>.
<sup>b</sup> The PRHC concentrations were calculated based on depletion studies conducted in our laboratory or by others in HepaRG cells or in primary human hepatocytes <sup>5,7,13</sup>. Cortisol, PGH, and oxytocin are not depleted by hepatocytes and/or HepaRG cells and therefore their concentrations were not adjusted.

### 2.4 CRISPR-Cas9–mediated gene knockdown and cortisol treatment

Since the above studies showed that cortisol was the main driver of transporter expression in HepaRG cells, subsequent mechanistic studies were performed using only cortisol. Differentiated HepaRG cells plated on collagen I-coated plates were transfected with TrueGuide synthetic sgRNAs targeting *NR3C1* (GR), *NR1I3* (CAR), *NR1I2* (PXR), *HNF4A* (HNF4α), or *HNF1A* (HNF1α), or with a non-targeting sgRNA control (sgRNA-NC). sgRNAs were complexed with TrueCut Cas9 Protein v2 and delivered as ribonucleoprotein complexes using Lipofectamine CRISPRMAX in Opti-MEM I according to the manufacturer’s instructions. After 48 h of transfection, cells were returned to dexamethasone-free maintenance medium and treated with 1× cortisol or vehicle for 48 h, with media refreshed every 24 h. Editing at each target locus was verified by Sanger sequencing and cleavage analysis as described below. The same transfection and treatment scheme was used for transporter activity experiments performed in 96-well plates.

### 2.5 RNA isolation and quantitative real-time PCR (qPCR)

Total RNA was isolated using the PureLink RNA Mini Kit and treated with DNase I after elution from the spin columns. RNA concentration and purity were assessed using a NanoDrop One Microvolume UV-Vis spectrophotometer (Thermo Fisher Scientific, Waltham, MA, USA). cDNA was synthesized using the High-Capacity cDNA Reverse Transcription Kit. qPCR was performed on a QuantStudio 5 Real-Time PCR System (Thermo Fisher Scientific, Waltham, MA, USA) using TaqMan Fast Advanced Master Mix and the TaqMan Gene Expression Assays listed above. *GAPDH* served as the housekeeping gene. Technical triplicates were averaged for each independent experiment before statistical analysis. Relative mRNA expression was calculated using the 2^-ΔΔCt method ^8^. For hormone-treatment experiments, expression was normalized to the corresponding vehicle control. For CRISPR experiments, expression was normalized to sgRNA-NC + vehicle. Data with Ct ≥ 36 were excluded.

### 2.6 Verification of gene editing by Sanger sequencing and cleavage analysis

Genomic DNA (gDNA) was isolated from matched cell lysates from the above mRNA isolation (before DNase I treatments) using the Genomic DNA Clean & Concentrator-10 kit. Regions flanking each sgRNA cut site (∼400–600 bp) were amplified by PCR using target-specific primers listed in Supplementary Table 1. PCR products were confirmed by agarose gel electrophoresis and purified using the PureLink PCR Purification Kit. Purified amplicons were submitted for Sanger sequencing (GENEWIZ, South Plainfield, NJ, USA), and editing was verified qualitatively by inspection of the chromatograms. Editing was further assessed using the GeneArt Genomic Cleavage Detection Kit (a DNA sequence mismatch-sensitive endonuclease-based assay) according to the manufacturer’s instructions. Briefly, PCR amplicons spanning each target site were denatured and re-annealed to form heteroduplex DNA, digested with the mismatch-detection nuclease, and resolved by agarose gel electrophoresis. Agarose gel electrophoresis was conducted using PowerPac Basic Power Supply (Bio-Rad, Hercules, CA, USA) at 100 V for 2 to 3 hours depending on gel size. Gels were stained after electrophoresis with SYBR Gold Nucleic Acid Gel Stain for 15 min. Representative Sanger chromatograms and cleavage assay results are shown in Supplementary Figures 1–6.

### 2.7 Transporter activity assays

For transporter activity experiments, HepaRG cells plated on 96-well collagen I-coated plates were treated with 1× cortisol or vehicle for 48 h after CRISPR transfection, as indicated in the corresponding figures. Transporter-mediated uptake was measured for NTCP, OAT2, and OCT1 using established substrate/inhibitor pairs ^5,9^. At the end of treatment, cells were washed three times with warm HBSS++ or Na^+^-free HBSS (to determine NTCP activity), as appropriate, and pre-equilibrated in the same buffer for 15 min at 37°C. Na^+^-free HBSS contains 138 mM of choline chloride, 1.26 mM of calcium chloride, 0.49 mM of magnesium chloride, 0.4 mM of magnesium sulfate, 5.33 mM of potassium chloride, 0.44 mM of monopotassium phosphate, 4.17 mM of potassium bicarbonate, 0.34 mM of dipotassium phosphate, and 5.56 mM of D-Glucose. Uptake was initiated by replacing the pre-incubation buffer with substrate solution in the presence or absence of inhibitor (or in Na^+^ free buffer) and was allowed to proceed for 15 min at 37°C (determined in prior experiments to be within initial linear uptake range ^7^). For NTCP, uptake was measured using 50 nM [^3^H]TA in HBSS++, as well as in Na^+^-free HBSS. For OAT2 and OCT1, uptake was measured using 40 nM [^3^H]cGMP ± inhibitor (200 μM BSP) and 8.8 μM [^14^C]metformin ± inhibitor (200 μM pyrimethamine), respectively. Uptake was terminated by rapid aspiration followed by three washes with ice-cold sodium-free HBSS. Cells were lysed in 200 μL/well lysis buffer from the PureLink RNA Mini Kit. A 150-μL aliquot was mixed with 4 mL Ecoscint ORIGINAL and radioactivity was quantified using a Tri-Carb B3110TR liquid scintillation counter (PerkinElmer, Waltham, MA, USA). The remaining lysate was reserved for RNA isolation and protein quantification. Net transporter-mediated uptake was calculated by subtracting uptake measured under inhibitor conditions from total uptake.

### 2.8 Protein extraction and quantification for transporter activity normalization

Total protein was quantified from the same samples used for transporter activity. After liquid scintillation counting, the 50-μL lysate reserved for RNA isolation was applied to the RNA spin column, and the first flow-through was collected for protein recovery. Four volumes of ice-cold methanol were added to the flow-through, followed by incubation at −20°C for 2 hours. Samples were centrifuged at 19,000 × g for 15 min at 4°C, the supernatant was removed, and the pellet was air-dried for approximately 5 min. Protein pellets were desalted using the Wessel-Flügge method ^10^ and resuspended in 50 μL of 2% SDS in 10 mM Tris-HCl (pH 7.5). Protein concentration was determined using the Pierce BCA Protein Assay Kit according to the manufacturer’s instructions and used to normalize uptake rates for the matched samples.

### 2.9 Statistical Analysis

Data were analyzed using GraphPad Prism 11.0.0 (GraphPad Software, La Jolla, CA, USA). Results are presented as mean ± SD of independent experiments, with technical triplicates averaged within each independent experiment before statistical analysis. For PRHC and individual-hormone mRNA experiments, one-way ANOVA followed by Dunnett’s multiple-comparison test versus vehicle was used. For CRISPR knockdown mRNA experiments, two-way ANOVA followed by Tukey’s multiple-comparison test was used. For matched transporter activity comparisons between sgRNA-NC + cortisol and target sgRNA + cortisol groups, paired two-tailed Student’s *t* test was used. Statistical tests for each experiment are also indicated in the corresponding figure legends. A value of *P* < 0.05 was considered statistically significant.

## 3. Results

### 3.1 Among individual PRHs, only cortisol induces hepatic uptake NTCP, OAT2 and OCT1 mRNA expression in HepaRG cells

Compared with vehicle, at 1x T3 concentrations, cortisol significantly induced NTCP, OAT2 OCT1, OAT7, and OAT2B1 mRNA expression (Figure 1A, C, E, Supplementary Figure 7) but did not affect OATP1B1 mRNA expression. This induction was similar in magnitude to that by the PRHC. At 1× concentrations, progesterone, testosterone, placental growth hormone, the individual estrogens (E1–E4), and oxytocin did not significantly alter transporter mRNA expression. Although several treatments produced mean responses similar in magnitude of induction to those observed at 1× PRHC or cortisol, these effects did not reach statistical significance. At 10× T3 concentrations, the direction of the cortisol response was generally preserved, but only NTCP remained significantly increased relative to vehicle (Figure 1B, D, and F).

**Figure 1.**
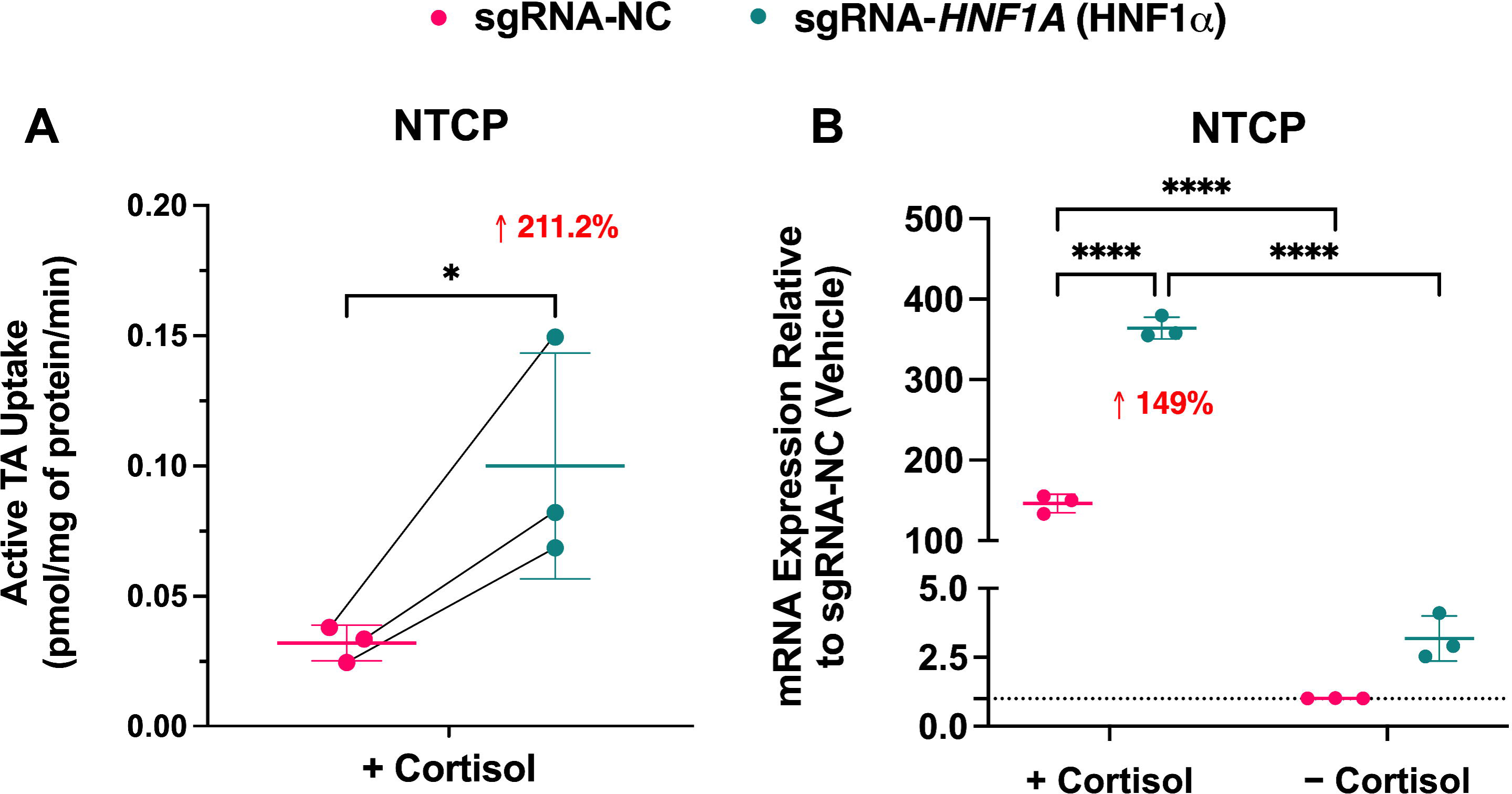
PRHC- and cortisol-mediated induction of NTCP, OAT2, and OCT1 mRNA expression in HepaRG cells. HepaRG cells were treated for 72 h with vehicle, a pregnancy-related hormone cocktail (PRHC), or individual hormones at 1× (A, C, E) or 10× (B, D, F) the geometric-mean third-trimester (T3) *in vivo* plasma concentrations (Table 1). NTCP (A, B), OAT2 (C, D), and OCT1 (E, F) mRNA expression, normalized to *GAPDH*, are expressed relative to vehicle (dotted line = 1.0). Where indicated, pooled estrogens are denoted E1+E2+E3+E4. At 1× T3 concentrations, PRHC and cortisol significantly increased NTCP and OCT1 mRNA expression to a similar extent, whereas only cortisol significantly increased OAT2 mRNA expression. At 10× T3 concentrations, only NTCP mRNA expression remained significantly induced despite a generally preserved direction of response for cortisol across all three transporters. Data are presented as mean ± SD from 3–5 independent experiments, each conducted in triplicate. Statistical analysis was performed by one-way ANOVA followed by Dunnett’s multiple-comparison test versus vehicle (\**P* < 0.05, \*\*\*\**P* < 0.0001). Ctrl, vehicle control; C, cortisol; P, progesterone; T, testosterone; PGH, placental growth hormone; E1, estrone; E2, estradiol; E3, estriol; E4, estetrol; O, oxytocin.

### 3.2 Cortisol, but not other PRHs, alters the mRNA expression of regulatory factors in HepaRG cells

Compared with vehicle, PRHC and cortisol at 1× T3 concentrations significantly increased CAR, PXR, and HNF4α mRNA expression in HepaRG cells (Figure 2A, C, E). Other PRHs, including progesterone, testosterone, placental growth hormone, the individual estrogens (E1–E4), and oxytocin, did not significantly alter the expression of these regulatory factors at 1× T3. At 10× T3 concentrations, PXR mRNA expression remained significantly increased by PRHC and cortisol, whereas CAR mRNA expression was no longer significantly altered and HNF4α mRNA expression remained significantly increased only after cortisol treatment (Figure 2B, D, and F).

**Figure 2.**
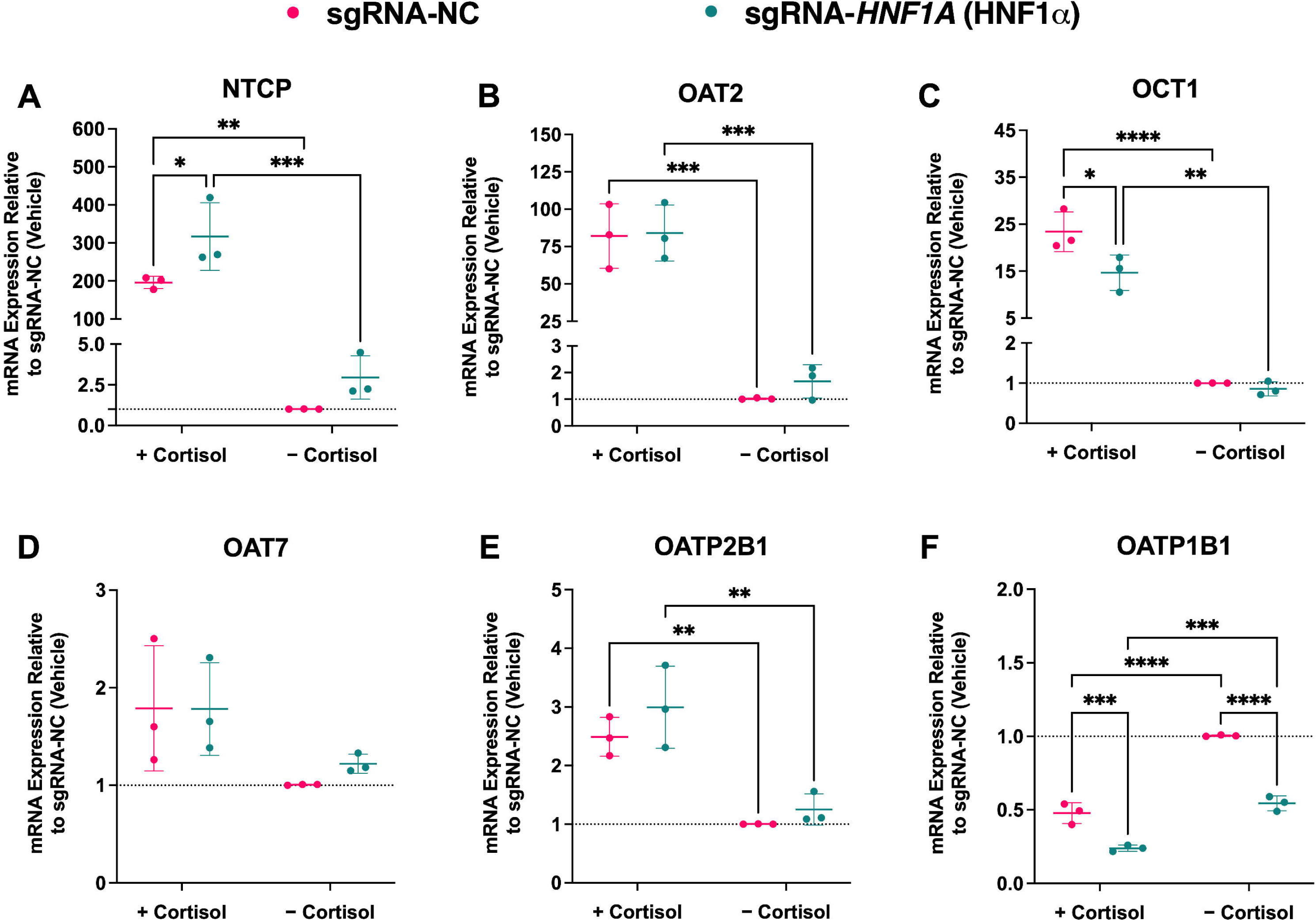
PRHC- and cortisol-mediated induction of PXR, CAR, and HNF4α mRNA expression in HepaRG cells. HepaRG cells were treated for 72 h with vehicle, PRHC, or individual hormones at 1× (A, C, E) or 10× (B, D, F) the geometric mean third-trimester (T3) *in vivo* plasma concentrations (Table 1). PXR (A, B), CAR (C, D), and HNF4α (E, F) mRNA expression, normalized to *GAPDH*, are expressed relative to vehicle (dotted line, 1.0). Where indicated, pooled estrogens are denoted E1+E2+E3+E4. At 1× T3 concentrations, PRHC and cortisol significantly increased PXR, CAR, and HNF4α mRNA expression to similar extents. At 10× T3 concentrations, PXR remained significantly increased by both PRHC and cortisol, whereas CAR was no longer significantly altered and HNF4α remained significantly increased only after cortisol treatment. Data are presented as mean ± SD from 3–5 independent experiments, each analyzed in triplicate. Statistical significance was assessed by one-way ANOVA followed by Dunnett’s multiple-comparison test versus vehicle (\**P* < 0.05, \*\**P* < 0.01, \*\*\**P* < 0.001, \*\*\*\**P* < 0.0001). Ctrl, vehicle control; C, cortisol; P, progesterone; T, testosterone; PGH, placental growth hormone; E1, estrone; E2, estradiol; E3, estriol; E4, estetrol; O, oxytocin.

At 1× T3 concentrations, PRHC and cortisol significantly increased FXR mRNA expression and decreased AhR mRNA expression, whereas GR mRNA expression was modestly decreased by PRHC and HNF1α mRNA expression was not significantly altered. In addition, 10× T3 PRHC and cortisol decreased HNF1α and AhR mRNA expression and modestly increased FXRα mRNA expression (Supplementary Figure 8). No other individual hormone or pooled estrogens (E1+E2+E3+E4) significantly altered PXR, CAR, or HNF4α mRNA expression at 10× concentrations.

### 3.3 GR knockdown attenuates cortisol-induced mRNA expression of NTCP, OAT2, and OCT1 in HepaRG cells

To determine whether cortisol-dependent transporter induction requires GR, *NR3C1* was knocked down by CRISPR-Cas9 in HepaRG cells. A preliminary 1× cortisol time-course showed that major transporter and regulatory-factor mRNA responses were already evident by 48 h and similar in magnitude to 72 hr (Supplementary Figure 9). Therefore, subsequent mechanistic knockdown studies were performed at 48 h to capture a strong cortisol response while maximizing our ability to measure a change in transporter activity which can lag change in trascript expression. After CRISPR-Cas9 transfection, cells were then treated with 1× T3 cortisol for 48 h. Editing of *NR3C1* in sgRNA-NR3C1-transfected cells was confirmed by Sanger sequencing and cleavage analysis (Supplementary Figures 1 and 6). In non-targeting sgRNA (sgRNA-negative control, NC)-transfected cells, as expected, cortisol significantly increased NTCP, OAT2, OCT1, and OATP2B1 mRNA expression and decreased OATP1B1 mRNA expression (Figure 3A–F). Specifically, NTCP, OAT2, OCT1, and OATP2B1 mRNA expression was induced by approximately 196-fold, 82-fold, 23-fold, and 2.5-fold, respectively, whereas OATP1B1 mRNA expression was reduced by 52% relative to sgRNA-NC + vehicle. GR knockdown markedly attenuated the cortisol response. Relative to sgRNA-NC + cortisol, mRNA induction significantly decreased to approximately 47-fold for NTCP, 38-fold for OAT2, and 9-fold for OCT1, corresponding to reductions of 76%, 54%, and 61%, respectively (Figure 3A–C). In this knockdown experiment, cortisol did not significantly increase OAT7 mRNA expression in sgRNA-NC-transfected cells (Figure 3D), unlike the modest induction observed in the earlier hormone-treatment experiments (Supplementary Figure 7). Although cortisol significantly increased OAT7 mRNA expression in sgRNA-*NR3C1*-transfected cells, OAT7 mRNA expression was not significantly different between sgRNA-NC + cortisol and sgRNA-*NR3C1* + cortisol groups (Figure 3D), indicating that GR knockdown did not significantly alter OAT7 mRNA expression under these conditions. Similarly, OATP2B1 mRNA induction was not significantly reduced after GR knockdown (Figure 3E). In contrast, cortisol-mediated repression of OATP1B1 mRNA expression persisted after GR knockdown, consistent with GR independent regulation under these conditions (Figure 3F). Basal transporter mRNA expression of all the above transporters in the absence of cortisol was comparable between sgRNA-NC- and sgRNA-*NR3C1*-transfected cells.

**Figure 3.**
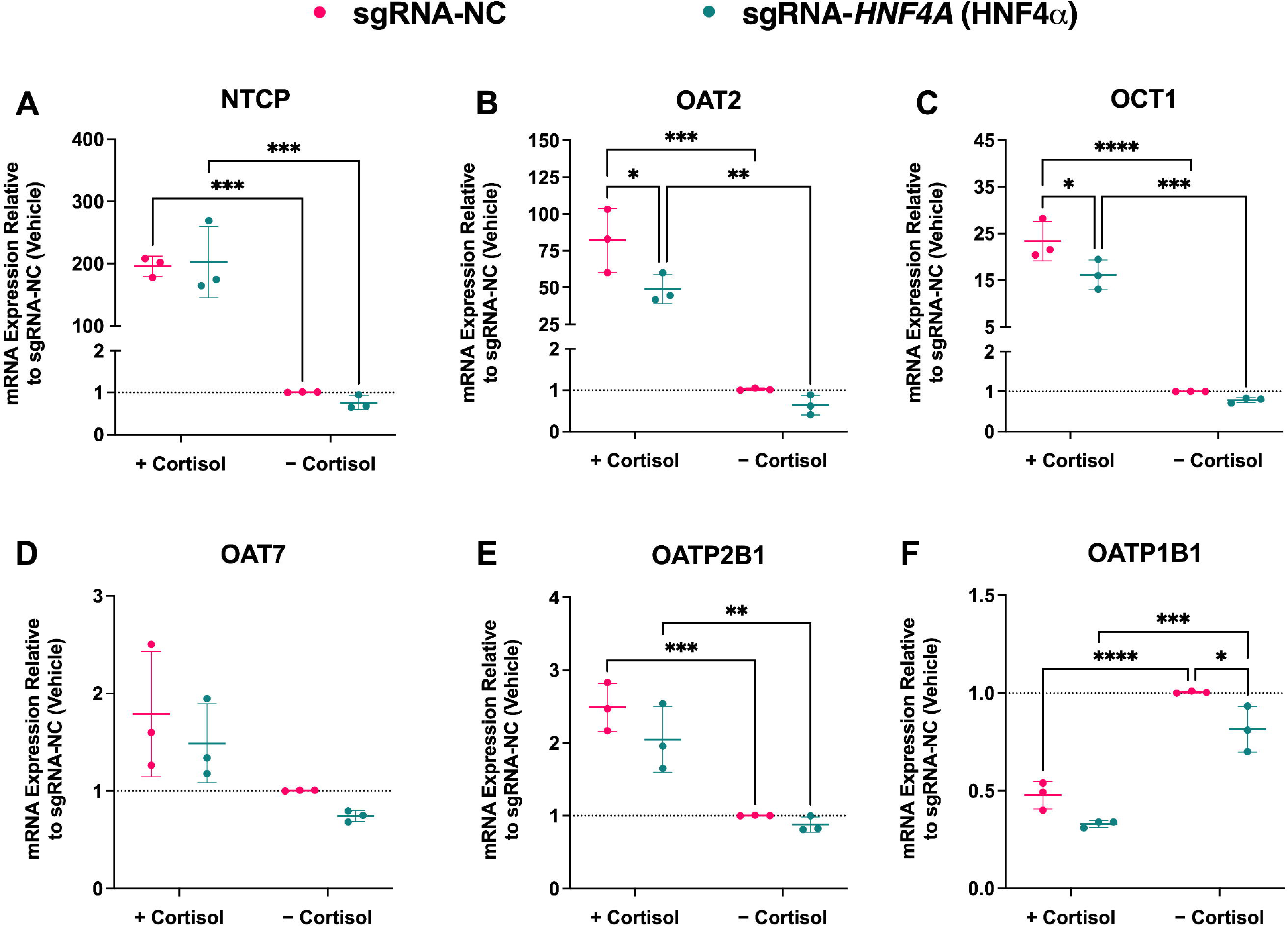
GR knockdown diminished cortisol-induced transporter mRNA expression in HepaRG cells. HepaRG cells were transfected with a non-targeting sgRNA control (sgRNA-NC) or GR-targeting sgRNA (sgRNA-*NR3C1*) and then treated with 1× T3 cortisol (Table 1) or vehicle for 48 h. NTCP (A), OAT2 (B), OCT1 (C), OAT7 (D), OATP2B1 (E), and OATP1B1 (F) mRNA expression, normalized to *GAPDH*, are expressed relative to sgRNA NC + vehicle (dotted line, 1.0). Points denote independent experiments; horizontal lines and error bars represent mean ± SD of n = 3 independent experiments, each conducted in triplicate. In sgRNA-NC-transfected cells, cortisol induced NTCP, OAT2, OCT1, and OATP2B1 mRNA expression and repressed OATP1B1 mRNA expression. GR knockdown significantly attenuated cortisol induction of NTCP, OAT2, and OCT1 mRNA expression by 76%, 54%, and 61%, respectively. GR knockdown did not significantly alter cortisol-mediated OATP2B1 and OAT7 mRNA induction and OATP1B1 mRNA repression. Statistical significance was assessed by two-way ANOVA followed by Tukey’s multiple comparison test (\**P* < 0.05; \*\**P* < 0.01; \*\*\**P* < 0.001; \*\*\*\**P* < 0.0001).

### 3.4 GR knockdown attenuates cortisol-induced NTCP, OAT2, and OCT1 activity and mRNA expression after 48 h of 1× T3 cortisol treatment

In sgRNA-NC-transfected cells, cortisol increased the active uptake of TA by NTCP, cGMP by OAT2, and metformin by OCT1 (Supplementary Figure 10). Quantitatively consistent with mRNA data, GR knockdown significantly attenuated these cortisol-induced transport activities by 72.8% for NTCP (Figure 4A), 53.2% for OAT2 (Figure 4B), and 60.0% for OCT1 (Figure 4C). In parallel, 48-h cortisol treatment robustly induced NTCP, OAT2, and OCT1 mRNA expression relative to matched sgRNA-NC + vehicle controls (Figure 4D–F). These mRNA inductions were significantly attenuated in sgRNA-*NR3C1*-transfected cells (NTCP: −75%; OAT2: −45%; OCT1: −66%), which recapitulates the mRNA findings shown in Figure 3 and aligned with the activity data (Figure 4A–C).

**Figure 4.**
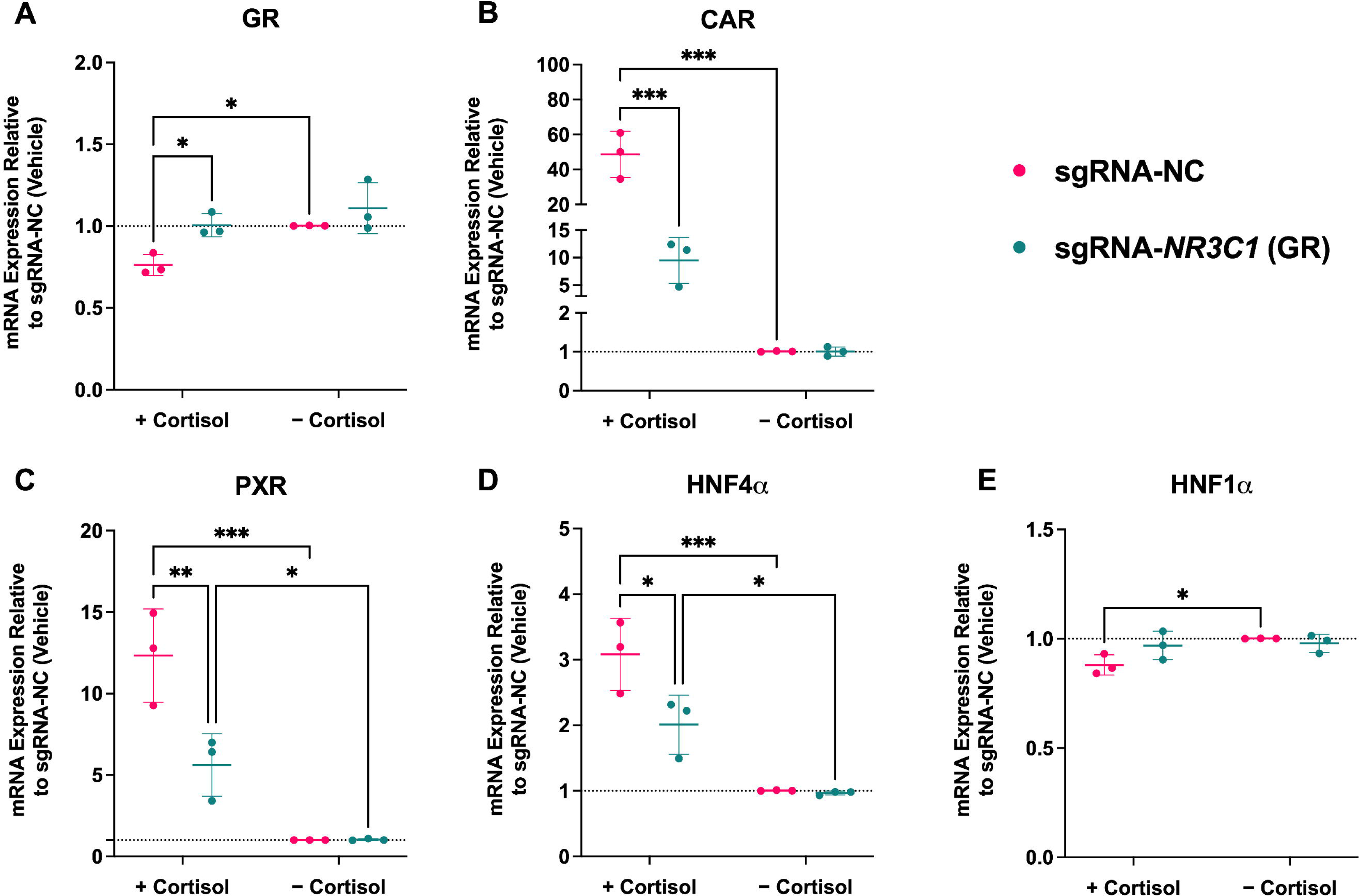
GR knockdown attenuates cortisol-induced NTCP, OAT2, and OCT1 activity and mRNA expression in HepaRG cells. HepaRG cells were transfected with sgRNA-NC or sgRNA-*NR3C1* and then treated with 1× T3 cortisol (Table 1) for 48 h. Top panels show mean active uptake (± SD) of TA ([^3^H]TA, 50 nM) by NTCP (A), cGMP ([^3^H]cGMP, 40 nM) by OAT2 (B), and metformin ([^14^C]metformin, 8.8 μM) by OCT1 (C). Passive uptake was determined using 50 nM [^3^H]TA with sodium-free HBSS, 40 nM [^3^H]cGMP with 200 μM BSP, and 8.8 μM [^14^C]metformin with 200 μM pyrimethamine, respectively. Bottom panels show NTCP (D), OAT2 (E), and OCT1 (F) mRNA expression normalized to *GAPDH* and expressed relative to sgRNA-NC + vehicle (dotted line, 1.0). In panels A–C, points denote independent experiments (each conducted in triplicate), and lines connect matched experimental replicates. For panels D–F, horizontal lines and error bars represent mean ± SD of *n* = 3 independent experiments, each conducted in triplicate. GR knockdown significantly reduced cortisol-induced NTCP, OAT2, and OCT1 activity and significantly attenuated the corresponding 48-h mRNA induction (NTCP: −75%; OAT2: −45%; OCT1: −66%). For transporter activity, statistical significance was assessed by paired two-tailed *t* test between sgRNA-NC + cortisol and sgRNA-*NR3C1* + cortisol (\**P* < 0.05, \*\**P* < 0.01, \*\*\**P* < 0.001). For mRNA, statistical significance was assessed by two-way ANOVA followed by Tukey’s multiple-comparison test (\*\**P* < 0.01, \*\*\**P* < 0.001, \*\*\*\**P* < 0.0001).

### 3.5 GR knockdown attenuates cortisol-induced CAR, PXR, and HNF4α mRNA expression and prevents cortisol-mediated decreases in GR and HNF1α mRNA in HepaRG cells

In sgRNA-NC-transfected HepaRG cells, 1× T3 cortisol significantly increased CAR, PXR, and HNF4α mRNA expression, while modestly decreasing GR and HNF1α mRNA expression (Figure 5A–E). Specifically, CAR, PXR, and HNF4α mRNA expression increased by approximately 48-fold, 12-fold, and 3-fold, respectively, whereas GR and HNF1α mRNA expression decreased by 24% and 12% relative to sgRNA-NC + vehicle. GR knockdown attenuated the cortisol response, reducing induction to approximately 8.5-fold for CAR (−82%), 6-fold for PXR (−50%), and 2-fold (−33%) for HNF4α. In addition, the cortisol-mediated decreases in GR and HNF1α mRNA observed in sgRNA-NC-transfected cells were no longer evident in sgRNA-*NR3C1*-transfected cells. In exploratory analyses of additional regulatory factors involved in hepatic transporter regulation, cortisol significantly increased FXR (∼3.8-fold), C/EBPβ (∼2.3-fold), and PGC-1α (∼3-fold) mRNA expression and decreased SHP (∼95%) and LRH-1 (∼62%) mRNA expression in sgRNA-NC-transfected HepaRG cells (Supplementary Figure 11A). Each of these cortisol-mediated changes was significantly attenuated by GR knockdown. Specifically, after GR knockdown, FXR, C/EBPβ, and PGC-1α mRNA expression was ∼2.31-fold, ∼1.41-fold, and ∼1.61-fold of control after cortisol treatment, respectively, while SHP and LRH-1 mRNA expression decreased by a lesser extent, 53% and 36%, respectively.

**Figure 5.**
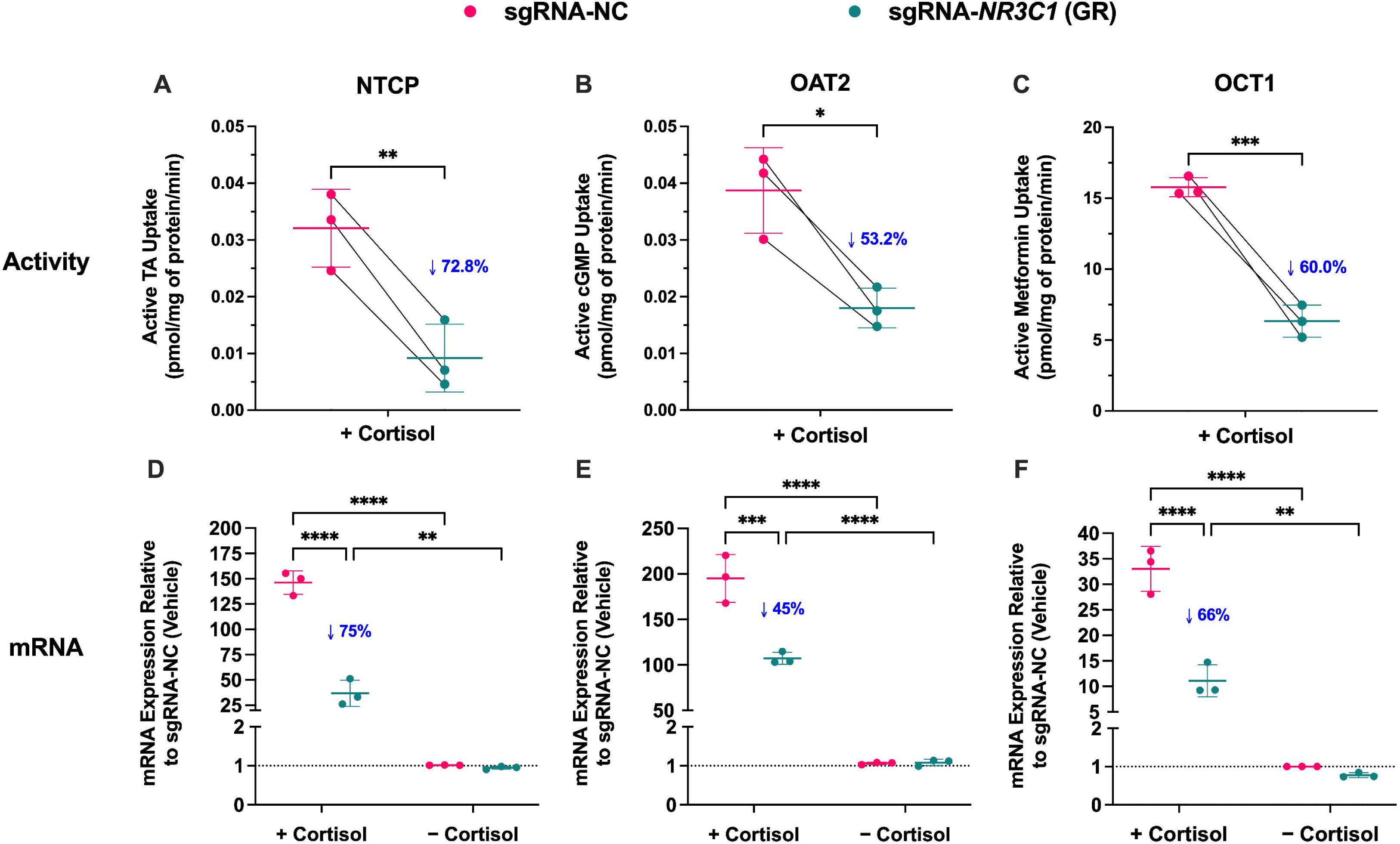
GR knockdown attenuates cortisol-induced CAR, PXR, and HNF4α mRNA expression and prevents cortisol-mediated decreases in GR and HNF1α mRNA in HepaRG cells. HepaRG cells were transfected with sgRNA-NC or sgRNA-*NR3C1* and then treated with 1× T3 cortisol (Table 1) or vehicle for 48 h. GR (A), CAR (B), PXR (C), HNF4α (D), and HNF1α (E) mRNA expression, normalized to *GAPDH*, are expressed relative to sgRNA-NC + vehicle (dotted line, 1.0). Points denote independent experiments. Horizontal lines and error bars represent mean ± SD of n = 3 independent experiments, each conducted in triplicate. In sgRNA-NC-transfected cells, cortisol increased CAR, PXR, and HNF4α mRNA expression and decreased GR and HNF1α mRNA expression. GR knockdown attenuated cortisol induction of CAR (−82%), PXR (−50%), and HNF4α (−33%) mRNA and prevented the cortisol-mediated decreases in GR and HNF1α mRNA. Statistical significance was assessed by two-way ANOVA followed by Tukey’s multiple-comparison test (\**P* < 0.05, \*\**P* < 0.01, \*\*\**P* < 0.001).

### 3.6 Individual CAR or PXR knockdown does not significantly alter cortisol-mediated changes in hepatic transporter or regulatory factor mRNA expression in HepaRG cells

To assess whether CAR (*NR1I3*) or PXR (*NR1I2*) contributes to cortisol-mediated transcriptional regulation in HepaRG cells, CAR or PXR was knocked down by CRISPR-Cas9 and cells were treated with 1× T3 cortisol for 48 h. Editing of *NR1I3* and *NR1I2* in sgRNA-*NR1I3*- and sgRNA-*NR1I2*-transfected cells was confirmed by Sanger sequencing and cleavage analysis (Supplementary Figures 2, 3, and 6). Cortisol-mediated changes in transporter mRNA expression (induction of NTCP, OAT2, OCT1, and OATP2B1 mRNA and repression of OATP1B1 mRNA) were preserved in both sgRNA- *NR1I3*- and sgRNA-*NR1I2*-transfected cells (Supplementary Figures 12 and 13). Transporter mRNA expression in cortisol-treated knockdown cells did not differ significantly from that in cortisol-treated sgRNA-NC cells. Similarly, in these experiments, cortisol-mediated mRNA changes in regulatory factors (decreased GR mRNA and increase CAR, PXR, and HNF4α mRNA) were preserved between sgRNA-NC and sgRNA- *NR1I3*- or sgRNA-*NR1I2*-transfected cells (Supplementary Figures 14 and 15). Together, these data indicate that individual CAR or PXR knockdown does not significantly alter cortisol-mediated changes in transporter or regulatory factor mRNA expression under these conditions. Exploratory analyses of additional regulatory factors further showed that this broader cortisol-responsive profile was largely preserved after individual CAR or PXR knockdown, although FXR induction was modestly enhanced after CAR knockdown (Supplementary Figure 11B, C).

### 3.7 HNF4α knockdown attenuates cortisol-induced OAT2 and OCT1 mRNA expression and reduces basal OATP1B1 mRNA expression in HepaRG cells

Similarly, to assess whether HNF4α (*HNF4A*) contributes to cortisol-mediated transcriptional regulation in HepaRG cells, HNF4α was knocked down by CRISPR-Cas9 and cells were treated with 1× T3 cortisol for 48 h. Editing of *HNF4A* in sgRNA-*HNF4A*-transfected cells was confirmed by Sanger sequencing and cleavage analysis (Supplementary Figures 4 and 6). HNF4α knockdown attenuated cortisol-induced OAT2 and OCT1 mRNA expression in HepaRG cells, reducing OAT2 induction from approximately 82-fold to 49-fold and OCT1 induction from 23-fold to 16-fold, corresponding to reductions of 40.2% and 30.0%, respectively (Figure 6B, C). In contrast, cortisol-mediated changes in NTCP, OAT7, and OATP2B1 mRNA expression were not significantly altered in sgRNA-*HNF4A*-transfected cells relative to sgRNA-NC-transfected cells (Figure 6A, D, E). OATP1B1 mRNA expression was reduced by approximately 22% in sgRNA-*HNF4A*-transfected cells under vehicle-treated conditions compared to sgRNA-NC-transfected cells (Figure 6F), indicating that HNF4α knockdown lowers basal OATP1B1 mRNA expression. mRNA expression of the regulatory factors (GR, CAR, PXR, HNF4α, and HNF1α) was also not significantly altered by HNF4α knockdown (Supplementary Figure 16). These data suggest that HNF4α contributes to cortisol-mediated induction of OAT2 and OCT1, but not to the regulation of the other transporters or regulatory factors examined. Exploratory analyses of other regulatory factors showed that HNF4α knockdown modestly attenuated cortisol-induced FXR mRNA expression, whereas cortisol-mediated changes in C/EBPβ, SHP, PGC-1α, LRH-1, and FOXA2 mRNA expression were not significantly altered (Supplementary Figure 11D).

### 3.8 HNF1α knockdown enhances cortisol-induced NTCP mRNA expression, attenuates OCT1 induction, and reduces basal OATP1B1 mRNA expression in HepaRG cells

Lastly, to assess whether HNF1α (*HNF1A*) contributes to cortisol-mediated transcriptional regulation in HepaRG cells, HNF1α was knocked down by CRISPR-Cas9 and cells were treated with 1× T3 cortisol for 48 h. Editing of *HNF1A* in sgRNA-*HNF1A*-transfected cells was confirmed by Sanger sequencing and cleavage analysis (Supplementary Figure 5 and 6). HNF1α knockdown enhanced cortisol-induced NTCP mRNA expression and attenuated OCT1 mRNA induction in HepaRG cells. Specifically, NTCP induction increased from approximately 196-fold to 317-fold (+62%), whereas OCT1 induction decreased from 23-fold to 15-fold (−35%) in cortisol-treated sgRNA-HNF1A-transfected cells relative to cortisol-treated sgRNA-NC cells (Figure 7A, C). OATP1B1 mRNA expression was also reduced by approximately 50% in sgRNA-*HNF1A*-transfected cells under both vehicle- and cortisol-treated conditions compared to sgRNA-NC-transfected cells (Figure 7F), indicating that HNF1α knockdown lowers basal OATP1B1 mRNA expression rather than specifically enhancing cortisol-mediated repression. In contrast, cortisol-mediated changes in OAT2, OAT7, and OATP2B1 mRNA expression were not significantly altered by HNF1α knockdown (Figure 7B, D, and E). Among the regulatory factors examined, only GR mRNA expression was further decreased modestly (∼13% further decrease) in cortisol-treated sgRNA-*HNF1A*-transfected cells, whereas CAR, PXR, HNF4α, and HNF1α mRNA expression were not significantly altered relative to cortisol-treated sgRNA-NC cells (Supplementary Figure 17). Exploratory analyses of other regulatory factors showed that HNF1α knockdown modestly attenuated cortisol-induced FXR mRNA expression and modestly enhanced PGC-1α mRNA induction, whereas cortisol-mediated changes in C/EBPβ, SHP, LRH-1, and FOXA2 mRNA expression were not significantly altered (Supplementary Figure 11E).

**Figure 6.**
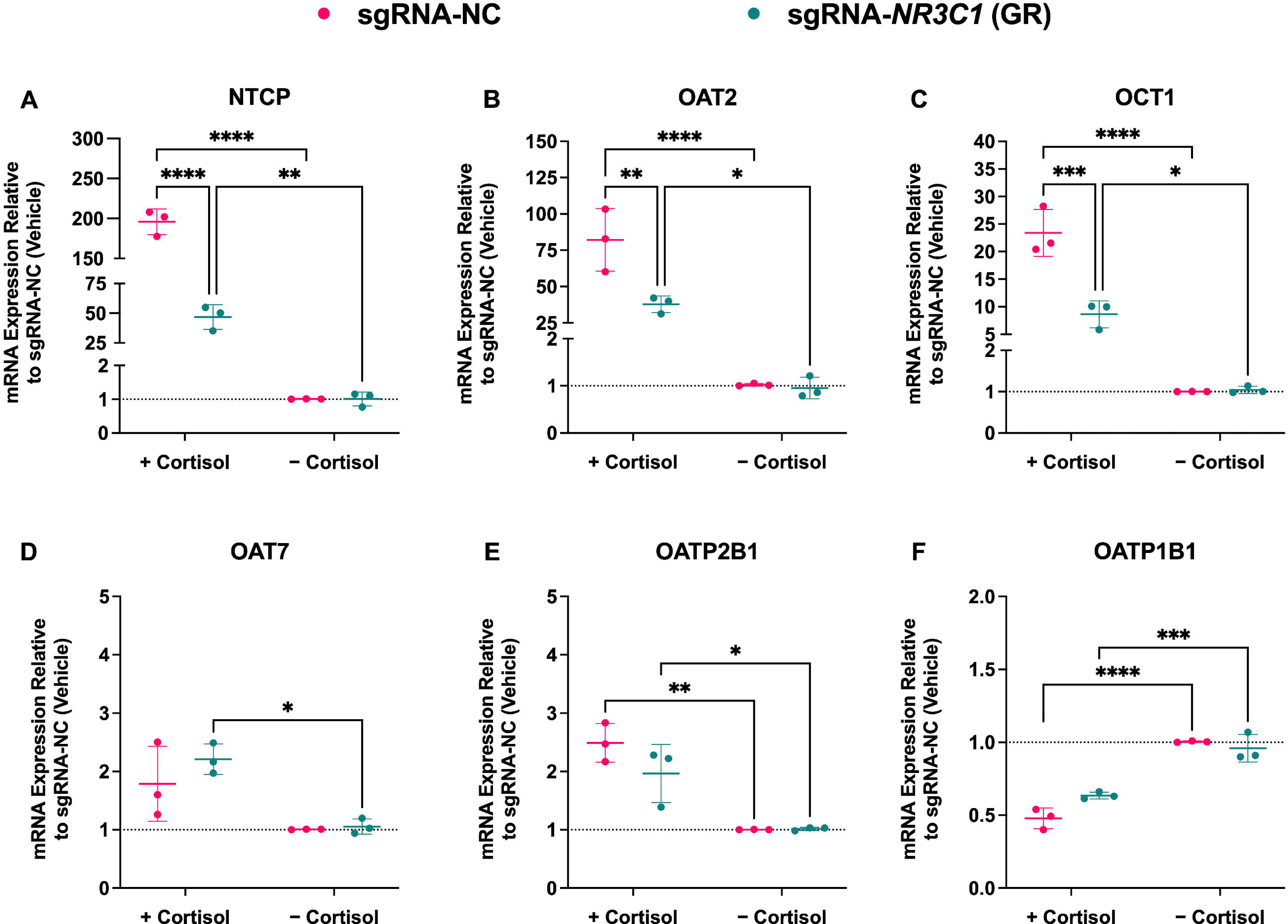
HNF4α knockdown attenuates cortisol-induced OAT2 and OCT1 mRNA expression in HepaRG cells. HepaRG cells were transfected with sgRNA-NC or sgRNA-*HNF4A* and then treated with 1× T3 cortisol (Table 1) or vehicle for 48 h. NTCP (A), OAT2 (B), OCT1 (C), OAT7 (D), OATP2B1 (E), and OATP1B1 (F) mRNA expression, normalized to *GAPDH*, are expressed relative to sgRNA-NC + vehicle (dotted line, 1.0). Points denote independent experiments. Horizontal lines and error bars represent mean ± SD of *n* = 3 independent experiments, each conducted in triplicate. HNF4α knockdown significantly attenuated cortisol induction of OAT2 (−40%) and OCT1 (−30%) mRNA but did not significantly alter cortisol-mediated changes in NTCP, OAT7, or OATP2B1 mRNA expression. Additionally, HNF4α knockdown significantly reduced OATP1B1 mRNA expression under vehicle-treated conditions, indicating reduced basal OATP1B1 expression. Statistical significance was determined using two-way ANOVA with Tukey’s post hoc test (\**P* < 0.05; ** *P* < 0.01; *** *P* < 0.001; \*\*\*\**P* < 0.0001).

**Figure 7.**
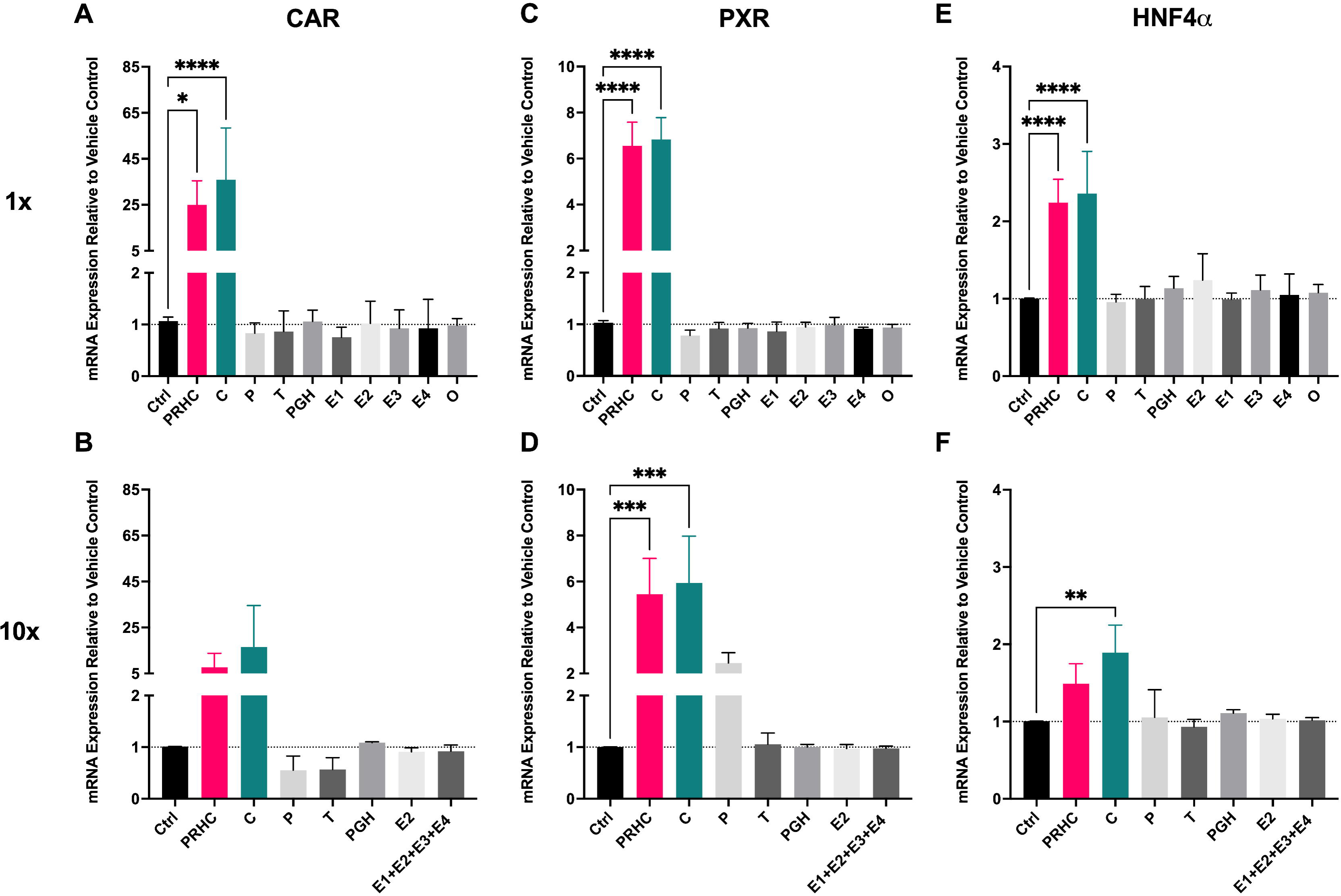
HNF1α knockdown differentially modulates cortisol-induced transporter mRNA expression in HepaRG cells. HepaRG cells were transfected with sgRNA-NC or sgRNA-*HNF1A* and then treated with 1× T3 cortisol (Table 1) or vehicle for 48 h. NTCP (A), OAT2 (B), OCT1 (C), OAT7 (D), OATP2B1 (E), and OATP1B1 (F) mRNA expression, normalized to *GAPDH*, are expressed relative to sgRNA-NC + vehicle (dotted line, 1.0). Points denote independent experiments. Horizontal lines and error bars represent mean ± SD of *n* = 3 independent experiments, each conducted in triplicate. HNF1α knockdown enhanced cortisol-induced NTCP mRNA expression (+62%) and attenuated cortisol-induced OCT1 mRNA expression (−35%), but did not significantly alter cortisol-mediated changes in OAT2, OAT7, or OATP2B1 mRNA expression. Additionally, HNF1α knockdown reduced OATP1B1 mRNA expression under both vehicle- and cortisol-treated conditions, indicating reduced basal OATP1B1 expression rather than a specific enhancement of cortisol-mediated repression. Statistical significance was assessed by two-way ANOVA followed by Tukey’s multiple-comparison test (\**P* < 0.05, \*\**P* < 0.01, \*\*\**P* < 0.001, \*\*\*\**P* < 0.0001).

### 3.9 HNF1α knockdown further enhances cortisol-induced NTCP activity and mRNA expression in HepaRG cells

HNF1α knockdown enhanced cortisol-induced NTCP activity in HepaRG cells after 48 h of 1× T3 cortisol treatment, increasing active TA uptake by 211.2% relative to cortisol-treated sgRNA-NC cells (Figure 8A). In parallel, NTCP mRNA expression was also significantly higher (+149%) in cortisol-treated sgRNA-HNF1A-transfected cells than in cortisol-treated sgRNA-NC-transfected cells (Figure 8B), consistent with the 48-h mRNA findings shown in Figure 7A.

**Figure 8.**
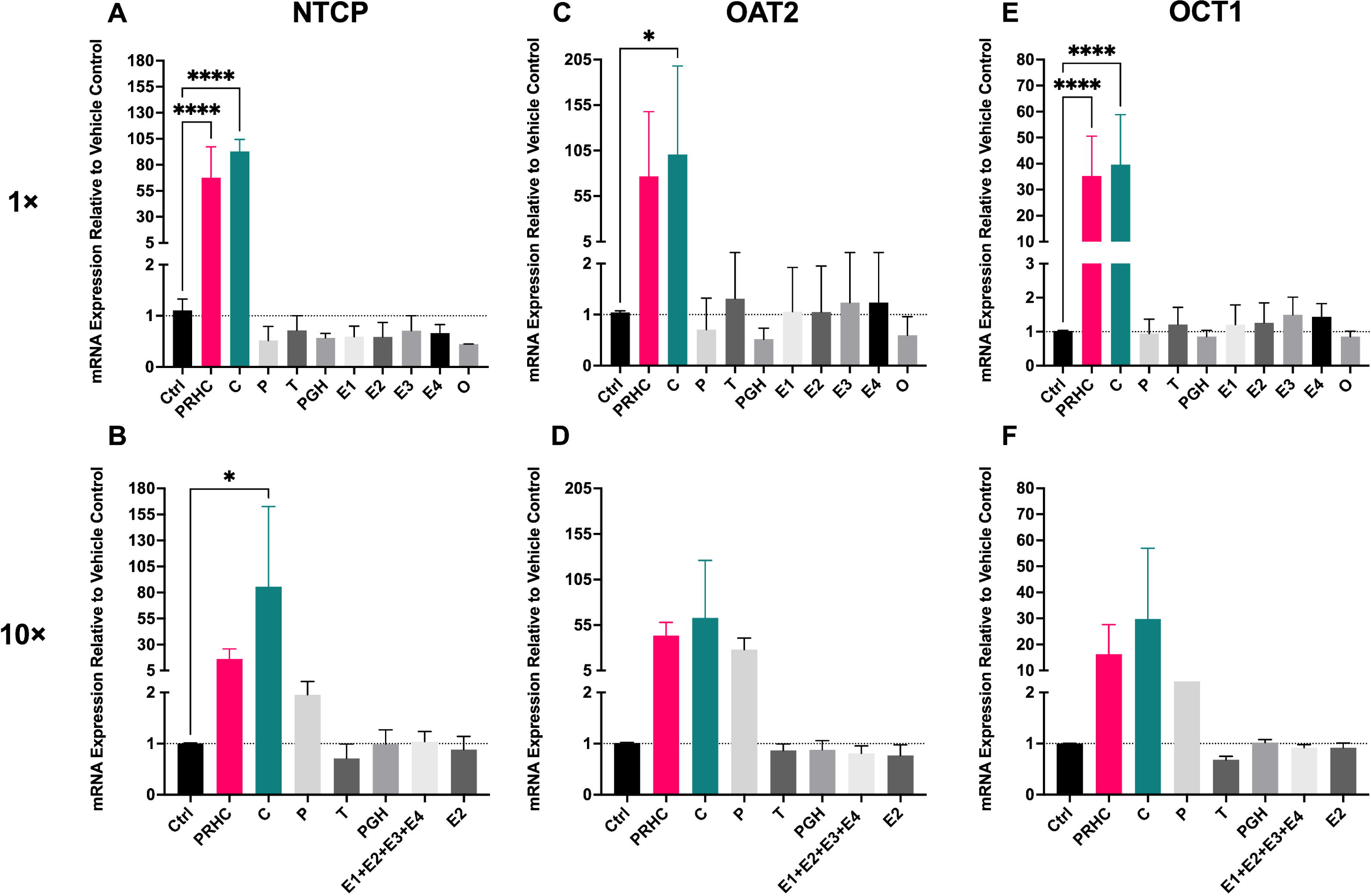
HNF1α knockdown enhanced cortisol-induced NTCP activity and mRNA expression in HepaRG cells. HepaRG cells were transfected with sgRNA-NC or sgRNA-*HNF1A* and then treated with 1× T3 cortisol (Table 1) or vehicle for 48 h. Panel A shows mean active uptake (± SD) of 50 nM [^3^H]TA by NTCP in cortisol-treated cells, determined with and without sodium-free HBSS. Panel B shows NTCP mRNA expression normalized to *GAPDH* and expressed relative to sgRNA-NC + vehicle (dotted line, 1.0). Points denote independent experiments. In panel A, lines connect matched experimental replicates. Horizontal lines and error bars represent mean ± SD of n = 3 independent experiments, each conducted in triplicate. HNF1α knockdown enhanced cortisol-induced NTCP activity (+211.2%) and NTCP mRNA expression (+149%) relative to sgRNA-NC-transfected cells. Statistical significance was assessed by paired two-tailed t test for transporter activity and by two-way ANOVA followed by Tukey’s multiple-comparison test for mRNA expression (P < 0.05, ***P < 0.0001).

These data indicate that HNF1α knockdown enhances cortisol-mediated NTCP induction at both the mRNA and activity levels.

### 3.10 Cortisol-regulated control genes show GR dependence and gene-specific effects of downstream factor knockdown

Because CYP3A4, CYP2B6, TAT, and UGT1A1 were previously identified as PRHC-responsive genes or hormone-responsive hepatic control genes (^7,11–13^, we evaluated whether the same transcription-factor knockdowns altered their response to cortisol in HepaRG cells (Supplementary Figures 18–22). In sgRNA-NC-transfected cells, cortisol induced CYP3A4 (151-fold), CYP2B6 (4-fold), and TAT (20-fold) mRNA expression (Supplementary Figures 18). GR knockdown markedly attenuated cortisol-induced CYP3A4 (−89%), CYP2B6 (−46%), and TAT (−67%) mRNA expression, consistent with a GR-dependent cortisol response (Supplementary Figure 18). HNF1α knockdown enhanced cortisol-induced TAT (+86%) and UGT1A1 (+81%) mRNA expression but did not blunt CYP3A4 and CYP2B6 induction (Supplementary Figure 19). HNF4α knockdown selectively reduced cortisol-induced CYP3A4 mRNA expression (−38%) (Supplementary Figure 20). In contrast, individual knockdown of CAR or PXR did not significantly alter the cortisol-mediated induction of these control genes (Supplementary Figures 21 and 22). These data show that cortisol also regulates additional hepatic control genes in a GR-dependent manner, but that downstream effects of HNF4α and HNF1α are gene-specific and distinct from the transporter response.

## Discussion

To our knowledge, this is the first integrated mechanistic study of PRH regulation of hepatic uptake transporters that involves identification of the perpetrator hormone, as well as targeted transcription factor knockdown using CRISPR-Cas9 and transporter activity readouts in HepaRG cells. This study identified cortisol as the principal pregnancy-related hormone driving hepatic uptake transporter regulation in HepaRG cells and defined the downstream transcriptional factors that shape this response. Among the individual hormones tested, cortisol reproduced most of the transporter and regulatory factor changes observed with the PRHC, whereas the other constituent hormones had little or no effect (Figure 1). At physiologic T3 plasma concentrations, cortisol significantly increased NTCP, OAT2, and OCT1 mRNA expression, as well as OAT7 and OATP2B1, and induced the regulatory factors PXR, CAR, and HNF4α (Figures 1 and 2, Supplementary Figure 7). By contrast, supraphysiologic (10×) exposure generally preserved the direction of the cortisol response, but only NTCP remained significantly induced under those conditions. These findings indicate that the T3-associated increase in plasma cortisol is sufficient to account for most of the PRHC response in this model and that the response is most clearly resolved at physiologic rather than supraphysiologic concentrations. The *in vivo* consequences and significance of these findings have been discussed in our previous manuscript ^7^.

The knockdown studies place GR as the main driver of cortisol-mediated transporter and regulatory factor regulation. In parallel mRNA and activity knockdown experiments, GR knockdown produced parallel decrease in cortisol-induced NTCP, OAT2, and OCT1 mRNA expression and transporter activity (Figure 4). This quantitative agreement between mRNA and activity strengthens the conclusion that cortisol regulates these transporters through a functionally relevant GR-dependent pathway rather than through isolated transcriptional effects. This interpretation is consistent with direct evidence that the human NTCP promoter is glucocorticoid responsive ^14^ and with prior work establishing OAT2 as an HNF4α-regulated gene (discussed below) ^15,16^. Taken together, the present data indicate that NTCP, OAT2, and OCT1 form the core cortisol-responsive uptake transporter module in HepaRG cells.

Compared with NTCP, OAT2, and OCT1, the cortisol-mediated regulation of OAT7, OATP2B1, and OATP1B1 was weaker and less consistent across experimental settings. In the 72-h hormone-treatment experiments, OAT7 and OATP2B1 mRNA expression was significantly increased by PRHC and cortisol, whereas OATP1B1 mRNA was not significantly altered (Supplementary Figure 7). In the 48-h GR knockdown experiments, cortisol increased OATP2B1 and decreased OATP1B1 mRNA expression in sgRNA-NC-transfected cells, but these responses were not significantly altered by GR knockdown (Figure 3). OAT7 mRNA induction also was not consistently detected in the cortisol-treated sgRNA-NC condition. Thus, compared with NTCP, OAT2, and OCT1, the regulation of OAT7, OATP2B1, and OATP1B1 by cortisol appears to be smaller in magnitude and more sensitive to experimental context. OATP1B3 was not included in the present hormone-treatment analysis because our previous study showed no change in OATP1B3 mRNA expression after PRHC treatment of HepaRG cells ^7^. Together, these results suggest that GR-mediated signaling is not the sole pathway of all hepatic uptake transporter responses previously observed with 1x T3 PRHC and cortisol and that some transporter endpoints may depend on additional transcriptional pathways or are more susceptible to variation introduced by treatment duration. Interestingly, the modest effect of PRH observed here and before ^5,7^ on OATP1B1 mRNA expression and activity in HepaRG cells or human hepatocytes is consistent with the modest reduction in oral clearance of RSV (an OATP1B1 substrate) in pregnant women ^4^.

GR knockdown also altered the regulatory factor response to cortisol. In sgRNA-NC-transfected cells, cortisol increased CAR, PXR, and HNF4α mRNA expression and modestly decreased GR and HNF1α mRNA expression (Figure 5). All these changes were significantly blunted or abolished after GR knockdown. Although basal GR mRNA expression was not decreased in vehicle-treated sgRNA-*NR3C1 (GR)*-transfected cells (Figure 5A), this likely reflects the location of the qPCR assay relative to the CRISPR-Cas9 target site. The sgRNA targeted exon 2 of *NR3C1*, whereas the TaqMan assay used to quantify GR mRNA spans the exon 4–5 junction. Therefore, CRISPR-edited *NR3C1* transcripts could still be detected by qPCR if the downstream exon 4–5 region remained present. Thus, the absence of reduced basal GR mRNA does not necessarily indicate absence of GR knockdown. Consistent with functional disruption of GR signaling, sgRNA-*NR3C1* transfection attenuated cortisol-mediated regulation of downstream transcription factors and transporter endpoints. A similar consideration applies to the other targeted transcription factors, for which unchanged basal mRNA expression after sgRNA transfection does not necessarily exclude disruption of protein function or downstream regulatory activity (Supplementary Figures 14–17). Together, these result places CAR, PXR, and HNF4α downstream, rather than upstream, of GR in cortisol response. In exploratory analyses of additional regulatory factors, cortisol also significantly increased FXR, C/EBPβ, and PGC-1α mRNA and decreased SHP and LRH-1 mRNA (Supplementary Figure 11). Each of these changes was attenuated by GR knockdown. Because these additional factors were not functionally interrogated (e.g., by knockdown studies), they should be interpreted cautiously.

Nevertheless, they suggest that cortisol does not simply induce a small number of isolated hepatic regulators but instead triggers a broader GR-dependent hepatic transcriptional program. The pattern involving FXR, SHP, and LRH-1 is especially notable because it does not mirror a simple canonical FXR “’I* SHP “’I* LRH-1 bile acid feedback response ^17^. Rather, it suggests that cortisol regulates these factors in a GR-sensitive manner that may influence transporter regulation more broadly. The additional control-gene data support this interpretation (Supplementary Figures 18–22). Cortisol induced CYP3A4, CYP2B6, and TAT mRNA expression in sgRNA-NC-transfected cells, and GR knockdown markedly attenuated each response. Individual knockdown of CAR or PXR did not significantly blunt cortisol-mediated induction of CYP3A4, CYP2B6, or TAT mRNA. HNF4α knockdown selectively attenuated cortisol-induced CYP3A4 mRNA, whereas HNF1α knockdown enhanced cortisol-induced TAT and UGT1A1 mRNA. Thus, these control-gene data further support a broad GR-dependent cortisol response, while showing that, compared to GR, the downstream transcription-factor effects are more gene-specific rather than uniform across hepatic genes.

Despite clear induction of CAR and PXR by cortisol, individual knockdown of either receptor did not significantly alter cortisol-mediated transporter induction or the main regulatory factor changes. Prior studies have shown that glucocorticoid receptor agonists can increase PXR and CAR expression through functional glucocorticoid-responsive elements (GREs) ^18–22^.

Therefore, the induction of CAR and PXR observed here is consistent with their regulation as part of a GR-responsive hepatic transcriptional program. However, the knockdown results indicate that neither CAR nor PXR is individually required for the transporter responses measured here. This differs from prior studies in HepaRG cells showing that cortisol-mediated CYP3A induction involves GR-dependent signaling that engages PXR and suggests that the pathway governing transporter induction is not simply the CYP3A pathway applied to transporter genes ^12,13^. A reasonable interpretation is that CAR and PXR are induced as part of the broader hepatic response to cortisol but are not obligate mediators of NTCP, OAT2, or OCT1 regulation. While these data do not exclude partial redundancy between CAR and PXR or contributions masked by incomplete knockdown, they do suggest that individual knockdown of either receptor is insufficient to transcriptionally dysregulate hepatic uptake transporter mRNA expression in HepaRG cells.

HNF4α showed a more selective role in hepatic transporter regulation. HNF4α knockdown significantly attenuated cortisol-induced OAT2 and OCT1 mRNA expression, did not significantly alter cortisol-mediated changes in NTCP, OAT7, or OATP2B1, and reduced basal OATP1B1 mRNA expression under vehicle-treated conditions. This selective pattern is consistent with the literature. OAT2 is a known HNF4α target, and recent work has shown that HNF4α helps define liver-specific glucocorticoid signaling ^15,16,23^. The present data extend that concept by showing that HNF4α contributes to the cortisol response of OAT2 and OCT1, but is not a global mediator of all cortisol-responsive hepatic transporters. The basal OATP1B1 effect further suggests that HNF4α may contribute modestly to constitutive OATP1B1 expression in HepaRG cells, even though it was not required for cortisol-mediated OATP1B1 repression. Together, these findings support a transporter-specific downstream role for HNF4α within the broader GR response.

Importantly, HNF1α knockdown enhanced cortisol-induced NTCP mRNA expression, reduced OCT1 induction, lowered OATP1B1 mRNA expression under both vehicle- and cortisol-treated conditions, and further decreased GR mRNA expression (Figure 7 and Supplementary Figure 17). These findings indicate that HNF1α’s role is transporter specific. The NTCP result was the most unexpected. HNF1α knockdown increased NTCP induction by cortisol treatment from 196-fold to 317-fold and, importantly, also increased NTCP activity by 211.2% (Figures 7 and 8). These data suggest that HNF1α restrains the full magnitude of the cortisol-driven NTCP response in HepaRG cells. Interestingly, while HNF1α does not contain GREs within its own gene sequence, its motif is in close proximity to GREs ^24^. Therefore, whether these findings reflect direct repression of NTCP by HNF1α or indirect modulation through other hepatic transcriptional circuits will require future studies. For OCT1, the data are consistent with a prior study showing that HNF1α contributes to OCT1 promoter activity and normal expression ^25^. For OATP1B1, the present data support a role for HNF1α in maintaining basal expression, because OATP1B1 mRNA was lower in HNF1α-knockdown cells regardless of whether cortisol was present (Figure 7). This interpretation is more accurate than describing the effect as enhanced cortisol-mediated repression, as the magnitude of OATP1B1 mRNA downregulation is similar between cortisol-treated sgRNA-NC- and sgRNA-*HNF1A*-transfected HepaRG cells. This is consistent with a previous study showing that HNF1α acts as a positive transcriptional regulator of OATP1B1 ^26–28^. For GR, the additional decrease in its mRNA expression after HNF1α knockdown is also consistent with prior evidence that HNF1α contributes to normal hepatic GR expression ^24^.

Our study has several limitations that can be addressed by future studies. First, the study was performed in HepaRG cells, which are useful for mechanistic dissection but may not reproduce the full complexity of pregnancy-associated regulation *in vivo* or in human hepatocytes. Second, most endpoints were assessed at the mRNA level, with functional activity measurements limited to NTCP, OAT2, and OCT1 because the activity of these transporters was most responsive to PRHC or cortisol treatment. Third, the regulation of basal OATP1B1 mRNA expression by HNF1α (and to a lesser extent, HNF4α) warrants further investigation as OATP1B1 activity is poor in HepaRG cells (data not shown). Therefore, future studies should focus on the involvement of HNF1α and HNF4α in the regulation of hepatic OATP1B1 in an *in vitro* system that better preserves OATP1B1 activity. Fourth, replication of our studies with T1 and T2 plasma cortisol concentrations will provide additional insights into the cortisol concentration-response relationships. Fifth, the CRISPR experiments are knockdowns rather than knockouts, so smaller contributions of CAR, PXR, HNF4α, or HNF1α may have been underestimated. However, this limitation does not undermine the main conclusion that GR is the primary mediator of the core transporter response, because GR knockdown reduced cortisol-induced NTCP, OAT2, and OCT1 activity by 72.8%, 53.2%, and 60.0%, respectively, in parallel with reductions in the corresponding mRNA induction (Figures 3 and 4). Similarly, HNF1α knockdown increased cortisol-induced NTCP activity by 211.2% (Figure 8). Thus, although incomplete knockdown may limit detection of smaller effects, the activity data support the conclusion that GR is the primary mediator of NTCP, OAT2, and OCT1 induction, whereas HNF1α selectively restrains NTCP induction. Sixth, our study focused primarily on hepatic uptake transporters as our previous data showed that the efflux transporters are only modestly affected by PRHC ^7^. Lastly, the study does not distinguish direct promoter-level regulation from indirect transcriptional effects. Promoter-reporter assays, chromatin occupancy studies, or rescue experiments would be needed to define the direct regulatory pathways.

In summary, cortisol was identified as the principal pregnancy-related hormone driving hepatic NTCP, OAT2, OCT1, OAT7 and OATP2B1 (but not OATP1B1) regulation in HepaRG cells and largely reproduced the effects of the pregnancy-related hormone cocktail at physiologic T3 plasma concentrations in HepaRG cells. This response was mediated primarily through GR, as GR knockdown markedly attenuated cortisol-induced NTCP, OAT2, and OCT1 expression and activity (Figure 9). In contrast, individual knockdown of CAR or PXR did not significantly alter the transporter response. HNF4α and HNF1α showed selective downstream effects, with HNF4α contributing to OAT2 and OCT1 induction and HNF1α differentially regulating NTCP, OCT1, and, together with HNF4α to a lesser extent, regulating basal OATP1B1 expression.

**Figure 9.**
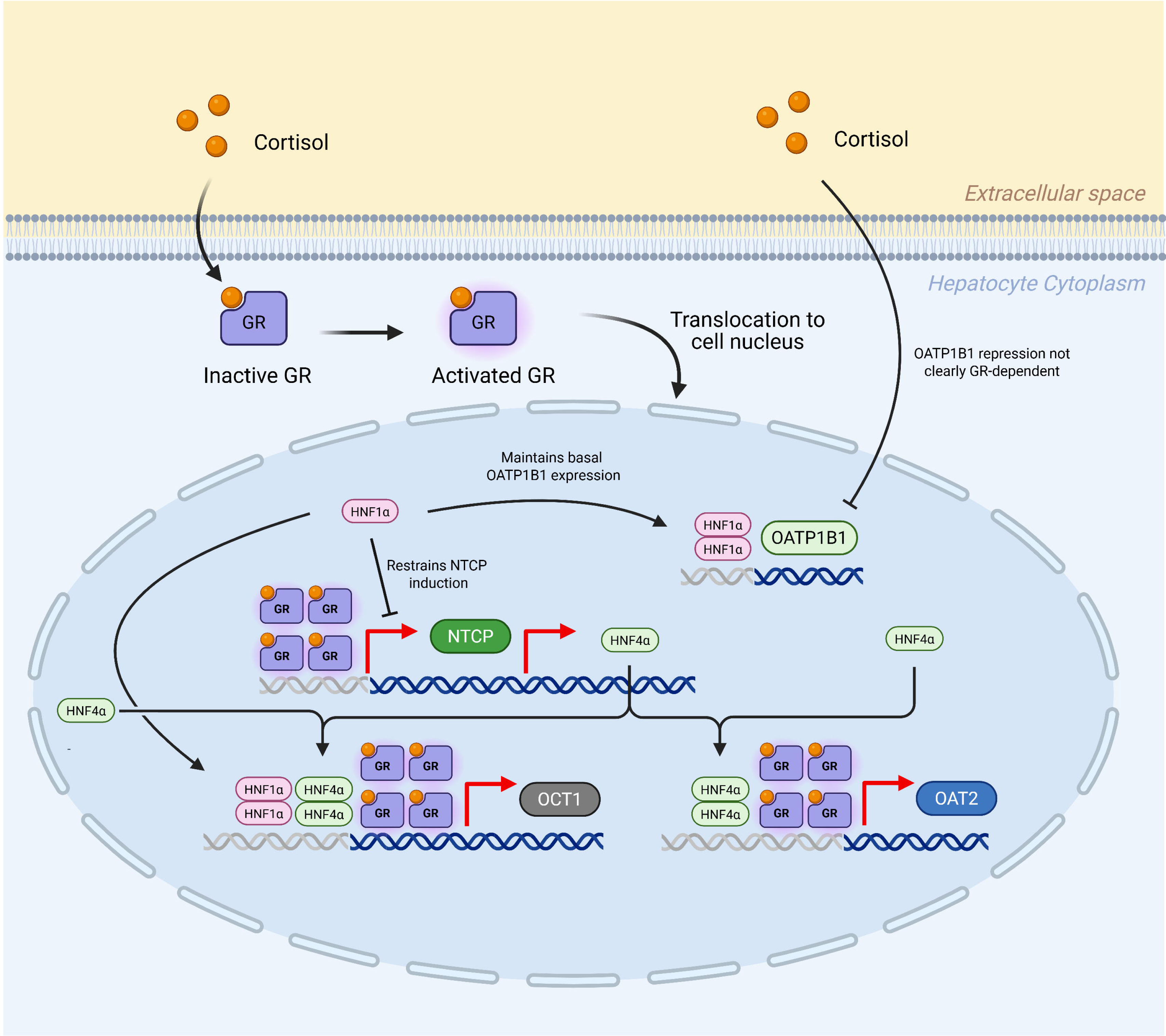
Proposed model of cortisol-mediated regulation of hepatic uptake transporters. Cortisol activates GR and promotes its nuclear translocation. Activated GR mediates the induction of NTCP, OAT2, and OCT1, as GR knockdown attenuated cortisol-induced mRNA expression and transporter activity for all three transporters. NTCP is shown as a direct GR-responsive transporter based on prior evidence of glucocorticoid responsiveness of the human NTCP promoter. HNF4α selectively contributes to cortisol-mediated OAT2 and OCT1 induction, as HNF4α knockdown attenuated the induction of these transporters but did not significantly alter NTCP induction. Thus, cortisol may regulate OAT2 and OCT1 in part indirectly through GR-dependent changes in HNF4α expression and/or HNF4α-dependent transcriptional activity. HNF1α differentially modulates transporter expression as HNF1α knockdown enhanced cortisol-mediated NTCP induction, attenuated cortisol-mediated OCT1 induction, and reduced basal OATP1B1 expression. Cortisol-mediated OATP1B1 repression is shown as GR-independent because GR knockdown did not reverse this effect. This model assumes a cortisol-GR centered pathway, with relatively modest cortisol-independent transporter modulation (e.g., OATP1B1) by HNF1α or HNF4α.

Together, these findings define a cortisol-GR-centered mechanism for pregnancy-associated hepatic uptake transporter regulation and provide a framework for improving prediction of transporter-based hepatic drug disposition during pregnancy. Our data also have implications for predicting hepatic uptake transporter changes after antenatal administration of corticosteroids to prevent neonatal respiratory distress syndrome due to premature birth ^29^.

## Acknowledgements

We thank Kai Wang (University of Washington) for his assistance with PCR, agarose gel electrophoresis, and sample preparation for Sanger sequencing.

## Data availability

The authors declare that all the data supporting the findings of this study are available within the paper and its Supplementary Data.

## CRediT authorship contribution statement

**Sejal Sharma:** Data curation, Formal analysis, Investigation, Methodology, Software, Validation, Visualization, Writing - original draft. **Yik Pui Tsang:** Data curation, Formal analysis, Investigation, Methodology, Software, Validation, Visualization, Writing - original draft. **Jashvant D. Unadkat:** Conceptualization, Funding acquisition, Methodology, Project administration, Resources, Supervision, Writing - review and editing.

## Footnotes

*-* This work was previously presented as a poster presentation at the following conference: Sharma S and Unadkat JD (2025) Regulation of expression and activity of hepatic transporters by pregnancy-related hormones in HepaRG cells. *14^th^ ISSX International Meeting*; 2025 Sep 21–24; Chicago, IL.
*-* Sejal Sharma and Yik Pui Tsang contributed equally to this work.
*- **Conflict of interest***: No author has an actual or perceived conflict of interest with the contents of this article.

## List of Abbreviations

AhR: (aryl hydrocarbon receptor)
ANOVA: (analysis of variance)
BCA: (bicinchoninic acid)
BSP: (sulfobromophthalein)
C/EBPβ: (CCAAT/enhancer-binding protein beta)
CAR: (constitutive androstane receptor)
Cas9: (CRISPR-associated protein 9)
cDNA: (complementary DNA)
cGMP: (cyclic guanosine monophosphate)
CRISPR: (clustered regularly interspaced short palindromic repeats)
CYP: (cytochrome P450)
DMSO: (dimethyl sulfoxide)
DNA: (deoxyribonucleic acid)
DPBS++: (Dulbecco’s phosphate-buffered saline with calcium and magnesium)
E1: (estrone)
E2: (estradiol)
E3: (estriol)
E4: (estetrol)
FXR: (farnesoid X receptor)
GAPDH: (glyceraldehyde-3-phosphate dehydrogenase)
gDNA: (genomic DNA)
GR: (glucocorticoid receptor)
GREs: (glucocorticoid-responsive elements)
HBSS: (Hanks’ balanced salt solution)
HNF1α: (hepatocyte nuclear factor 1 alpha)
HNF3β: (hepatocyte nuclear factor 3 beta)
HNF4α: (hepatocyte nuclear factor 4 alpha)
ITS-G: (insulin-transferrin-selenium)
LRH-1: (liver receptor homolog-1)
mRNA: (messenger RNA)
NC: (negative control)
NIH: (National Institutes of Health)
NTCP: (sodium/taurocholate cotransporting polypeptide)
OAT2: (organic anion transporter 2)
OAT7: (organic anion transporter 7)
OATP1B1: (organic anion transporting polypeptide 1B1)
OATP2B1: (organic anion transporting polypeptide 2B1)
OCT1: (organic cation transporter 1)
PBPK: (physiologically based pharmacokinetic)
PCR: (polymerase chain reaction)
PGC-1α: (peroxisome proliferator-activated receptor gamma coactivator 1-alpha)
PGH: (placental growth hormone)
PRH: (pregnancy-related hormone)
PRHC: (pregnancy-related hormone cocktail)
PXR: (pregnane X receptor)
qPCR: (quantitative real-time polymerase chain reaction)
RNA: (ribonucleic acid)
SD: (standard deviation)
SDS: (sodium dodecyl sulfate)
sgRNA: (single guide RNA)
sgRNA-NC: (non-targeting single guide RNA control)
SHP: (small heterodimer partner)
T3: (third trimester)
TA: (taurocholic acid)
UV-Vis: (ultraviolet-visible)

## References

1. Isoherranen N, Thummel KE. Drug metabolism and transport during pregnancy: how does drug disposition change during pregnancy and what are the mechanisms that cause such changes? Drug Metab Dispos. 2013;41(2):256–262. doi:10.1124/dmd.112.050245

2. Hebert M, Easterling T, Kirby B, et al. Effects of Pregnancy on CYP3A and P-glycoprotein Activities as Measured by Disposition of Midazolam and Digoxin: A University of Washington Specialized Center of Research Study. Clin Pharmacol Ther. 2008;84(2):248–253. doi:10.1038/clpt.2008.1

3. Unadkat JD, Wara DW, Hughes MD, et al. Pharmacokinetics and Safety of Indinavir in Human Immunodeficiency Virus-Infected Pregnant Women. Antimicrob Agents Chemother. 2007;51(2):783–786. doi:10.1128/AAC.00420-06

4. Pego ÁMG, Marques MP, Moreira F de L, et al. In Vivo Activity of Intestinal PßGlycoprotein and Hepatic Organic Anion Transporters Polypeptide in Pregnancy and Postpartum. The Journal of Clinical Pharmacology. 2025;65(1):7–17. doi:10.1002/jcph.6125

5. Benzi JR de L, Tsang YP, Unadkat JD. The effect of pregnancy-related hormones on hepatic transporters: studies with premenopausal human hepatocytes. Front Pharmacol. 2024;15. doi:10.3389/fphar.2024.1440010

6. Schock H, Zeleniuch-Jacquotte A, Lundin E, et al. Hormone concentrations throughout uncomplicated pregnancies: a longitudinal study. BMC Pregnancy Childbirth. 2016;16(1):146. doi:10.1186/s12884-016-0937-5

7. Sharma S, Unadkat JD. Regulation of expression and activity of hepatic transporters by pregnancy-related hormones in HepaRG cells. Drug Metabolism and Disposition. 2025;53(8):100118. doi:10.1016/j.dmd.2025.100118

8. Livak KJ, Schmittgen TD. Analysis of Relative Gene Expression Data Using Real-Time Quantitative PCR and the 2−ΔΔCT Method. Methods. 2001;25(4):402–408. doi:10.1006/meth.2001.1262

9. Hao T, Tsang YP, Yin M, Mao Q, Unadkat JD. Dysregulation of Human Hepatic Drug Transporters by Proinflammatory Cytokines. Journal of Pharmacology and Experimental Therapeutics. 2024;391(1):82–90. doi:10.1124/jpet.123.002019

10. Wessel D, Flügge UI. A method for the quantitative recovery of protein in dilute solution in the presence of detergents and lipids. Anal Biochem. 1984;138(1):141–143. doi:10.1016/0003-2697(84)90782-6

11. Khatri R, Fallon JK, Sykes C, et al. Pregnancy-Related Hormones Increase UGT1A1-Mediated Labetalol Metabolism in Human Hepatocytes. Front Pharmacol. 2021;12. doi:10.3389/fphar.2021.655320

12. Sachar M, Kelly EJ, Unadkat JD. Mechanisms of CYP3A Induction During Pregnancy: Studies in HepaRG Cells. AAPS J. 2019;21(3):45. doi:10.1208/s12248-019-0316-z

13. Zhang Z, Farooq M, Prasad B, Grepper S, Unadkat JD. Prediction of Gestational Age–Dependent Induction of In Vivo Hepatic CYP3A Activity Based on HepaRG Cells and Human Hepatocytes. Drug Metabolism and Disposition. 2015;43(6):836–842. doi:10.1124/dmd.114.062984

14. Eloranta JJ, Jung D, Kullak-Ublick GA. The Human Na+-Taurocholate Cotransporting Polypeptide Gene Is Activated by Glucocorticoid Receptor and Peroxisome Proliferator-Activated Receptor-γ Coactivator-1α, and Suppressed by Bile Acids via a Small Heterodimer Partner-Dependent Mechanism. Molecular Endocrinology. 2006;20(1):65–79. doi:10.1210/me.2005-0159

15. Yu Z, You G. Recent Advances on the Regulations of Organic Anion Transporters. Pharmaceutics. 2024;16(11):1355. doi:10.3390/pharmaceutics16111355

16. Popowski K, Eloranta JJ, Saborowski M, Fried M, Meier PJ, Kullak-Ublick GA. The Human Organic Anion Transporter 2 Gene Is Transactivated by Hepatocyte Nuclear Factor-4α and Suppressed by Bile Acids. Mol Pharmacol. 2005;67(5):1629–1638. doi:10.1124/mol.104.010223

17. Goodwin B, Jones SA, Price RR, et al. A Regulatory Cascade of the Nuclear Receptors FXR, SHP-1, and LRH-1 Represses Bile Acid Biosynthesis. Mol Cell. 2000;6(3):517–526. doi:10.1016/S1097-2765(00)00051-4

18. Pascussi J, Drocourt L, GerbalßChaloin S, Fabre J, Maurel P, Vilarem M. Dual effect of dexamethasone on CYP3A4 gene expression in human hepatocytes. Eur J Biochem. 2001;268(24):6346–6358. doi:10.1046/j.0014-2956.2001.02540.x

19. Pascussi JM, Drocourt L, Fabre JM, Maurel P, Vilarem MJ. Dexamethasone Induces Pregnane X Receptor and Retinoid X Receptor-α Expression in Human Hepatocytes: Synergistic Increase of CYP3A4 Induction by Pregnane X Receptor Activators. Mol Pharmacol. 2000;58(2):361–372. doi:10.1124/mol.58.2.361

20. Pascussi JM, Robert A, Moreau A, et al. Differential regulation of constitutive androstane receptor expression by hepatocyte nuclear factor4α isoforms†. Hepatology. 2007;45(5):1146–1153. doi:10.1002/hep.21592

21. Shi D, Yang D, Yan B. Dexamethasone transcriptionally increases the expression of the pregnane X receptor and synergistically enhances pyrethroid esfenvalerate in the induction of cytochrome P450 3A23. Biochem Pharmacol. 2010;80(8):1274–1283. doi:10.1016/j.bcp.2010.06.043

22. Daujat-Chavanieu M, Gerbal-Chaloin S. Regulation of CAR and PXR Expression in Health and Disease. Cells. 2020;9(11):2395. doi:10.3390/cells9112395

23. Hunter AL, Poolman TM, Kim D, et al. HNF4A modulates glucocorticoid action in the liver. Cell Rep. 2022;39(3):110697. doi:10.1016/j.celrep.2022.110697

24. Lin WY, Hu YJ, Lee YH. Hepatocyte nuclear factor-1α regulates glucocorticoid receptor expression to control postnatal body growth. American Journal of Physiology-Gastrointestinal and Liver Physiology. 2008;295(3):G542–G551. doi:10.1152/ajpgi.00081.2008

25. O’Brien VP, Bokelmann K, Ramírez J, et al. Hepatocyte Nuclear Factor 1 Regulates the Expression of the Organic Cation Transporter 1 via Binding to an Evolutionary Conserved Region in Intron 1 of the OCT1 Gene. J Pharmacol Exp Ther. 2013;347(1):181–192. doi:10.1124/jpet.113.206359

26. He YJ, Zhang W, Tu JH, et al. Hepatic Nuclear Factor 1α Inhibitor Ursodeoxycholic Acid Influences Pharmacokinetics of the Organic Anion Transporting Polypeptide 1B1 Substrate Rosuvastatin and Bilirubin. Drug Metabolism and Disposition. 2008;36(8):1453–1456. doi:10.1124/dmd.108.020503

27. Jung D, Hagenbuch B, Gresh L, Pontoglio M, Meier PJ, Kullak-Ublick GA. Characterization of the Human OATP-C (SLC21A6) Gene Promoter and Regulation of Liver-specific OATP Genes by Hepatocyte Nuclear Factor 1α. Journal of Biological Chemistry. 2001;276(40):37206–37214. doi:10.1074/jbc.M103988200

28. Furihata T, Satoh T, Yamamoto N, Kobayashi K, Chiba K. Hepatocyte Nuclear Factor 1 Alpha is a Factor Responsible for the Interindividual Variation of OATP1B1 mRNA Levels in Adult Japanese Livers. Pharm Res. 2007;24(12):2327–2332. doi:10.1007/s11095-007-9458-2

29. Anoshchenko O, Milad MA, Unadkat JD. Estimating fetal exposure to the Pßgp substrates, corticosteroids, by PBPK modeling to inform prevention of neonatal respiratory distress syndrome. CPT Pharmacometrics Syst Pharmacol. 2021;10(9):1057–1070. doi:10.1002/psp4.12674

