## Supplemental File for "Cortisol Drives Pregnancy-Associated Induction of Hepatic OAT2, NTCP, and OCT1 in HepaRG cells Through GR-, HNF1α-, and HNF4α-Dependent Signaling"

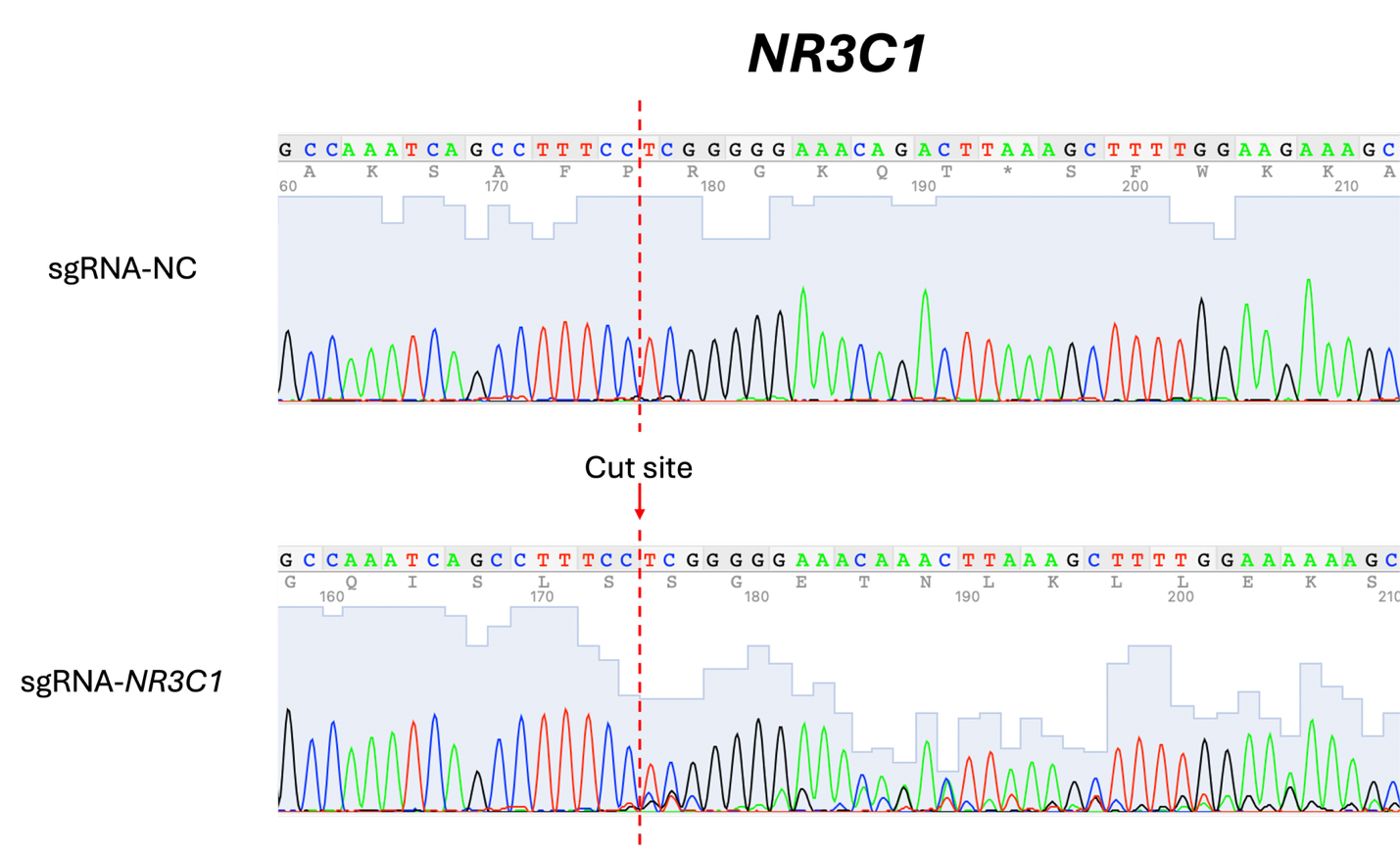

**Supplementary Figure 1. Representative Sanger sequencing chromatograms confirming CRISPR-Cas9 editing of *NR3C1* (GR) in HepaRG cells.** HepaRG cells were transfected with non-targeting sgRNA control (sgRNA-NC) or *NR3C1*-targeting sgRNA (sgRNA-*NR3C1*). Representative Sanger sequencing chromatograms of PCR amplicons spanning the *NR3C1* target site are shown. The dashed red line indicates the predicted Cas9 cut site. The sgRNA-NC chromatogram shows a clean sequence trace across the target region, whereas the sgRNA-*NR3C1* chromatogram shows mixed/overlapping peaks beginning at the cut site, consistent with CRISPR-Cas9-mediated editing and heterogeneous indel formation.

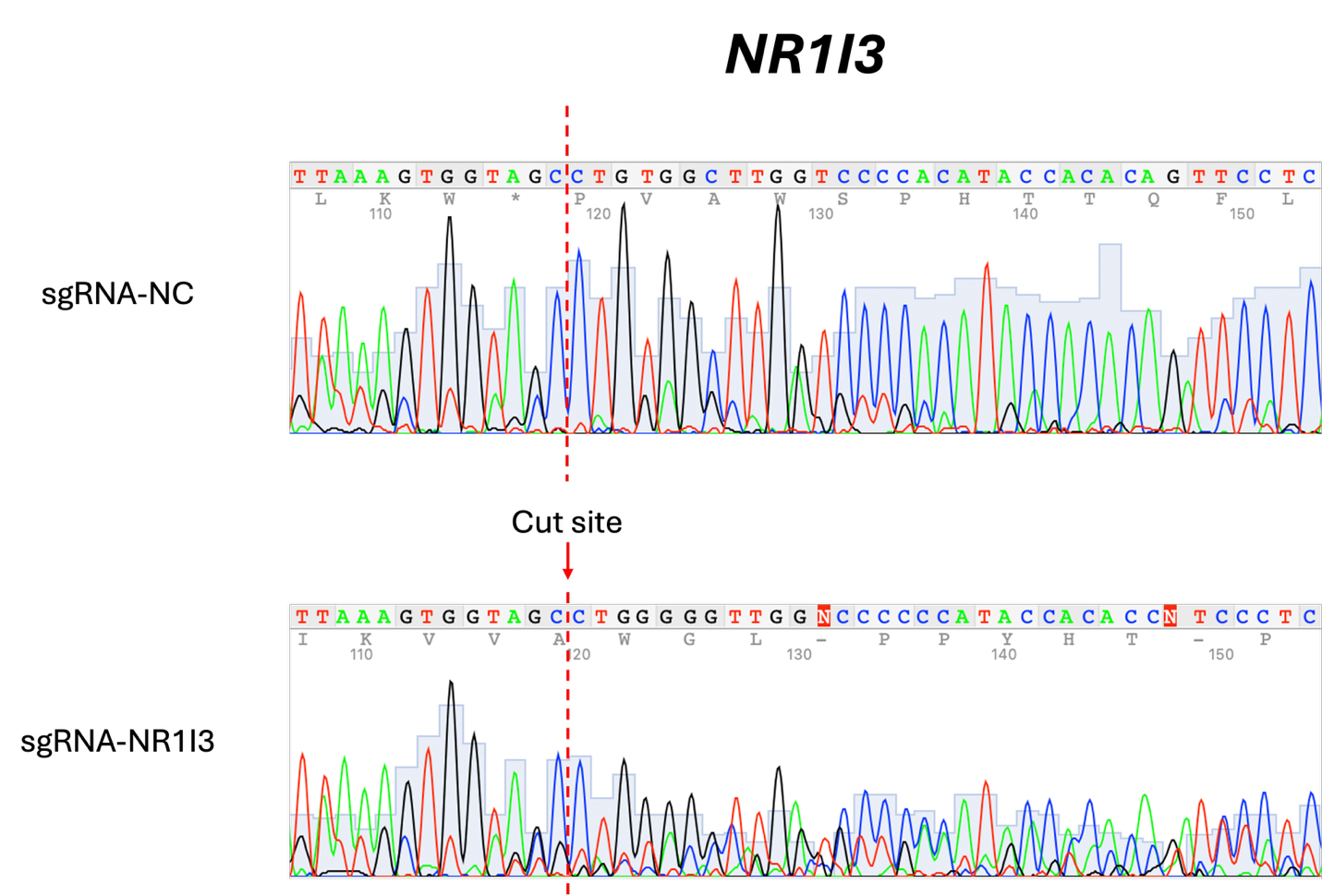
**Supplementary Figure 2. Representative Sanger sequencing chromatograms confirming CRISPR-Cas9 editing of *NR1I3* (CAR) in HepaRG cells.** HepaRG cells were transfected with non-targeting sgRNA control (sgRNA-NC) or *NR1I3*-targeting sgRNA (sgRNA-*NR1I3*). Representative Sanger sequencing chromatograms of PCR amplicons spanning the *NR1I3* target site are shown. The dashed red line indicates the predicted Cas9 cut site. The sgRNA-NC chromatogram shows a clean sequence trace across the target region, whereas the sgRNA-*NR1I3* chromatogram shows mixed/overlapping peaks beginning at the cut site, consistent with CRISPR-Cas9-mediated editing and heterogeneous indel formation.

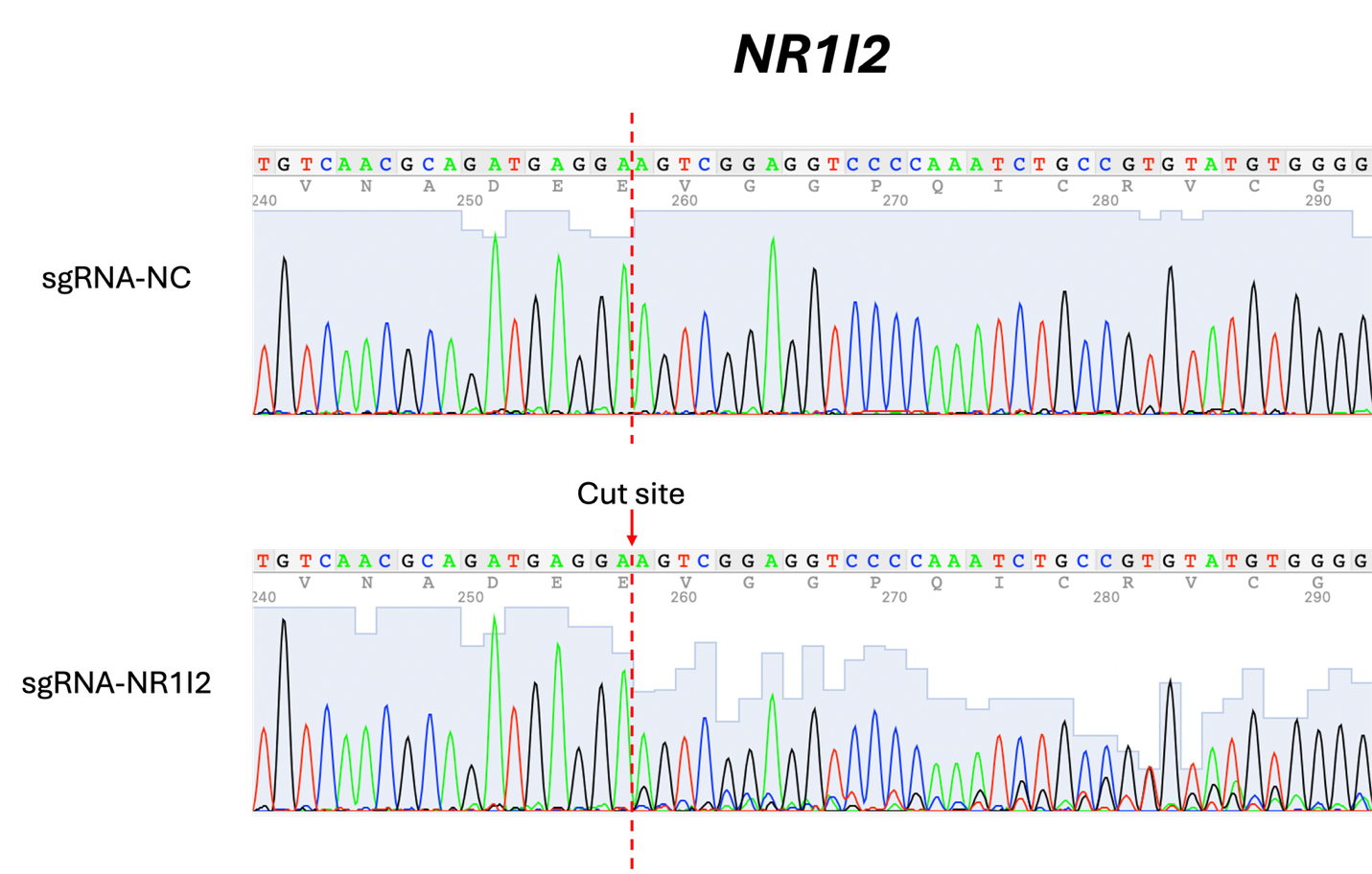

**Supplementary Figure 3. Representative Sanger sequencing chromatograms confirming CRISPR-Cas9 editing of *NR1I2* (PXR) in HepaRG cells.** HepaRG cells were transfected with non-targeting sgRNA control (sgRNA-NC) or *NR1I2*-targeting sgRNA (sgRNA-*NR1I2*). Representative Sanger sequencing chromatograms of PCR amplicons spanning the *NR1I2* target site are shown. The dashed red line indicates the predicted Cas9 cut site. The sgRNA-NC chromatogram shows a clean sequence trace across the target region, whereas the sgRNA-*NR1I2* chromatogram shows mixed/overlapping peaks beginning at the cut site, consistent with CRISPR-Cas9-mediated editing and heterogeneous indel formation.

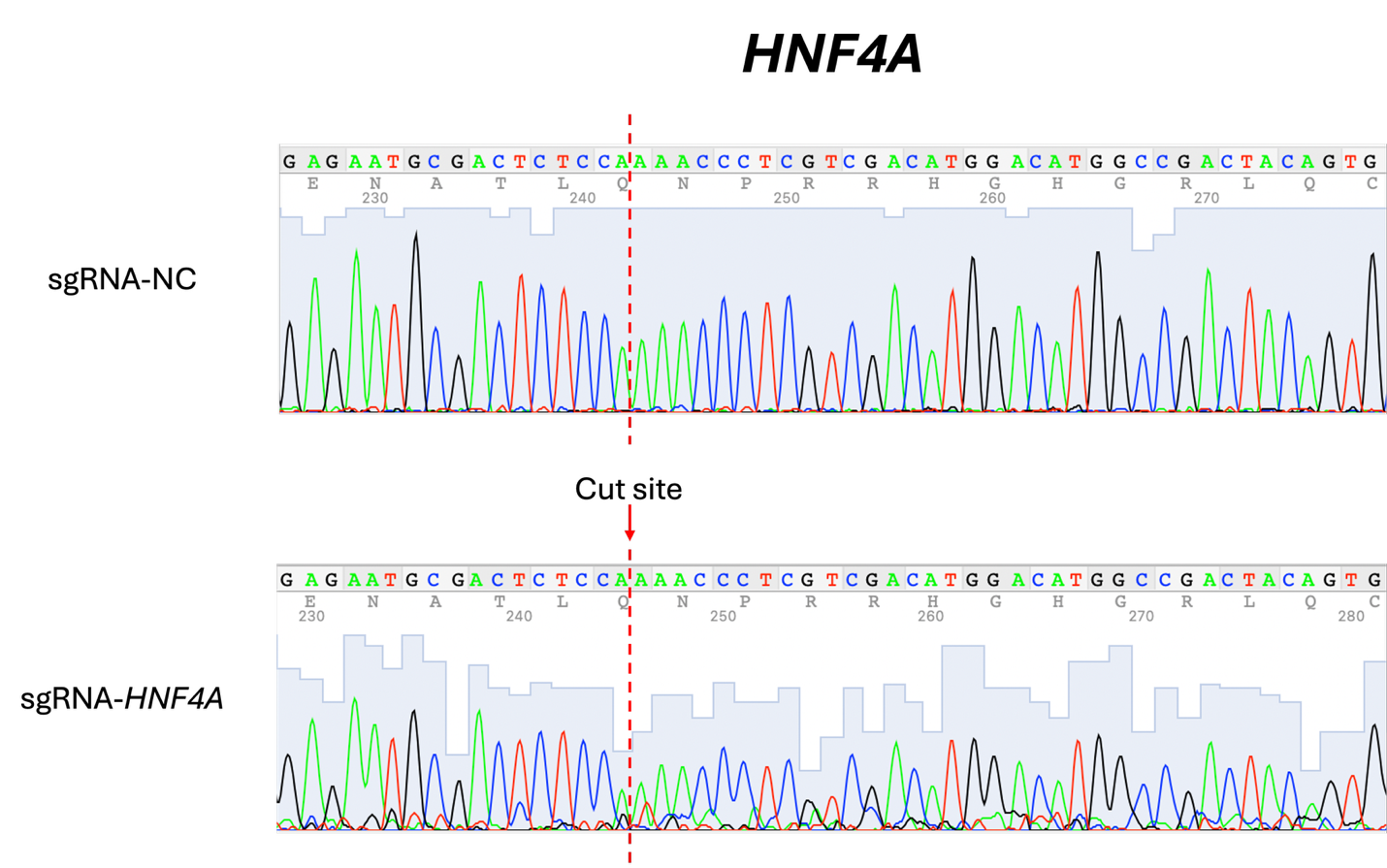

**Supplementary Figure 4. Representative Sanger sequencing chromatograms confirming CRISPR-Cas9 editing of *HNF4A* (HNF4α) in HepaRG cells.** HepaRG cells were transfected with non-targeting sgRNA control (sgRNA-NC) or *HNF4A*-targeting sgRNA (sgRNA-*HNF4A*). Representative Sanger sequencing chromatograms of PCR amplicons spanning the *HNF4A* target site are shown. The dashed red line indicates the predicted Cas9 cut site. The sgRNA-NC chromatogram shows a clean sequence trace across the target region, whereas the sgRNA-*HNF4A* chromatogram shows mixed/overlapping peaks beginning at the cut site, consistent with CRISPR-Cas9-mediated editing and heterogeneous indel formation.

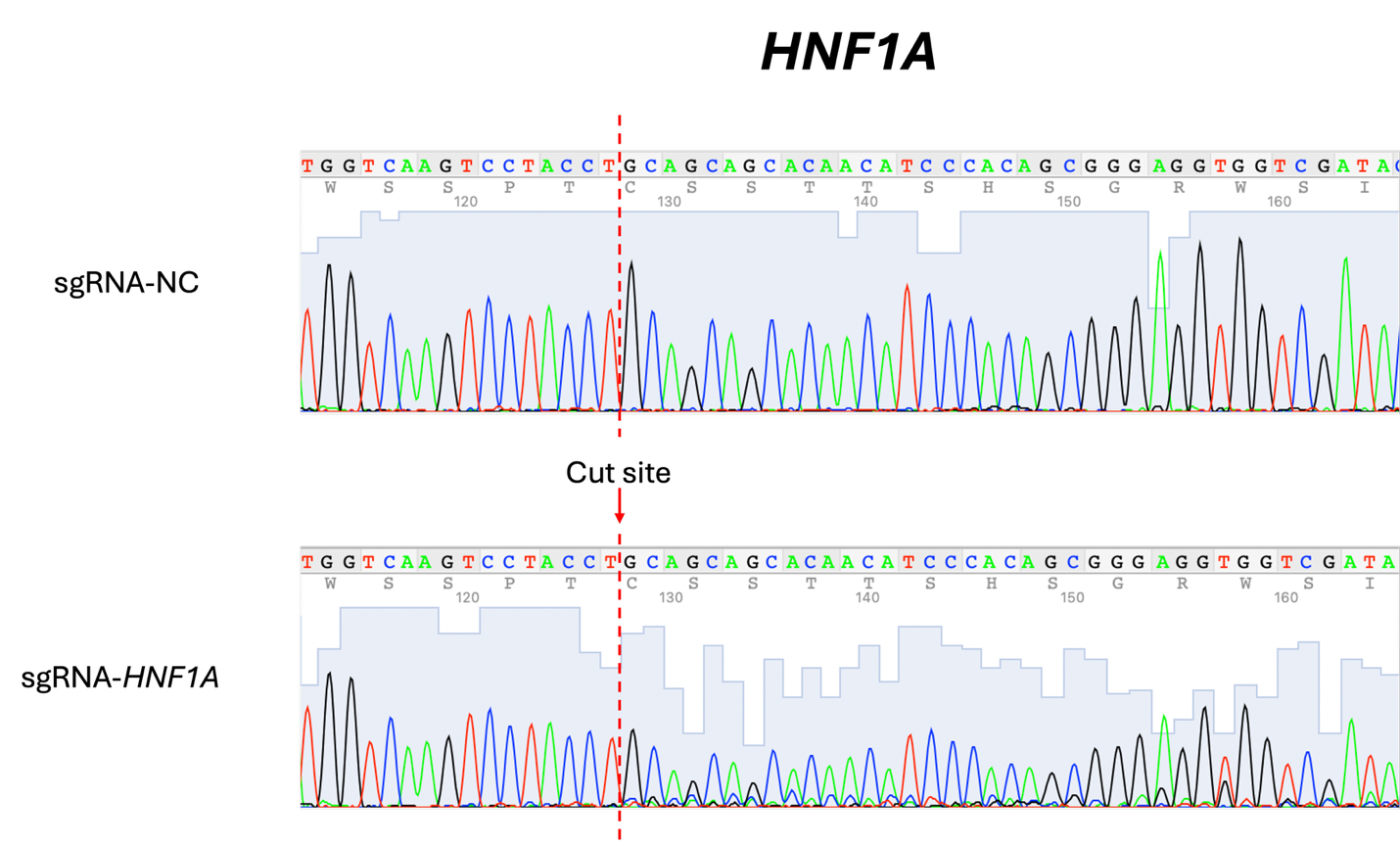

**Supplementary Figure 5. Representative Sanger sequencing chromatograms confirming CRISPR-Cas9 editing of *HNF1A* (HNF1α) in HepaRG cells.** HepaRG cells were transfected with non-targeting sgRNA control (sgRNA-NC) or *HNF1A*-targeting sgRNA (sgRNA-*HNF1A*). Representative Sanger sequencing chromatograms of PCR amplicons spanning the *HNF1A* target site are shown. The dashed red line indicates the predicted Cas9 cut site. The sgRNA-NC chromatogram shows a clean sequence trace across the target region, whereas the sgRNA-*HNF1A* chromatogram shows mixed/overlapping peaks beginning at the cut site, consistent with CRISPR-Cas9-mediated editing and heterogeneous indel formation.

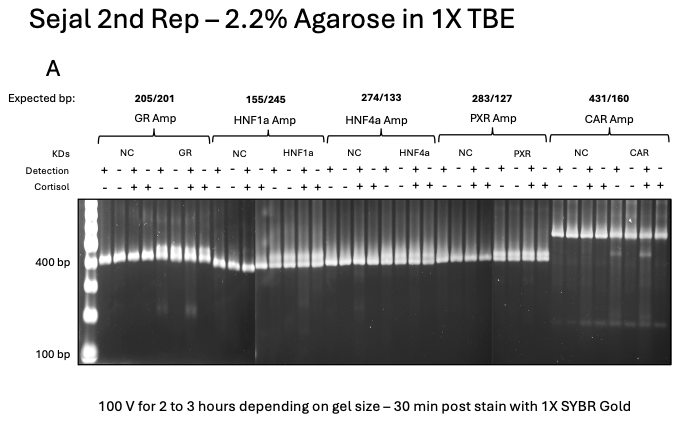

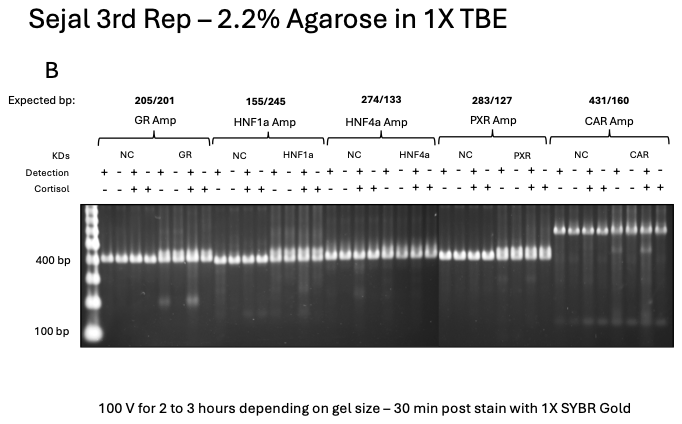

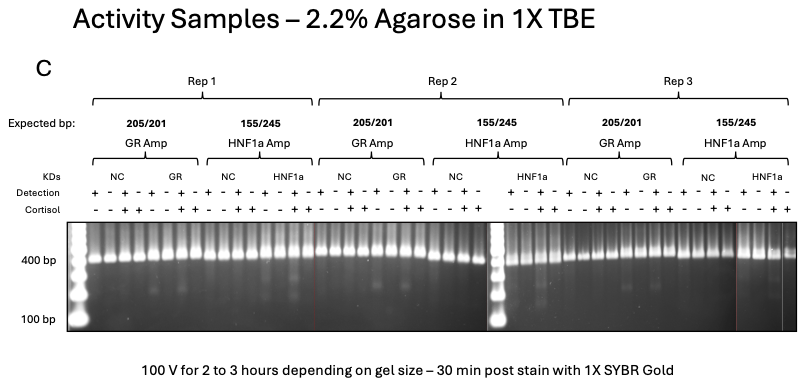

**Supplementary Figure 6. Cleavage analysis confirming CRISPR-Cas9 editing of *NR3C1, HNF1A, HNF4A, NR1I2,* and *NR1I3* in HepaRG cells.** (A, B) PCR amplicons spanning the *NR3C1, HNF1A, HNF4A, NR1I2,* and *NR1I3* target sites from replicate experiment 2 (A) and replicate experiment 3 (B) of the initial 48-h mRNA endpoint experiments. (C) PCR amplicons spanning the *NR3C1* and *HNF1A* target sites from three independent activity/parallel qPCR experiments, corresponding to the two knockdown conditions carried forward to functional activity studies. Amplicons from sgRNA-NC- and target sgRNA-transfected cells treated with vehicle or 1x cortisol were analyzed with (+) or without (−) mismatch-detection nuclease, as indicated. Expected cleavage fragment sizes (bp) are shown above each amplicon. Cleavage products of the expected sizes were predominantly observed in target sgRNA-transfected samples but not in sgRNA-NC controls, consistent with CRISPR-Cas9-mediated editing. Genomic DNA from replicate 1 of the initial 48-h mRNA endpoint experiments was not retained and therefore was unavailable for cleavage analysis.

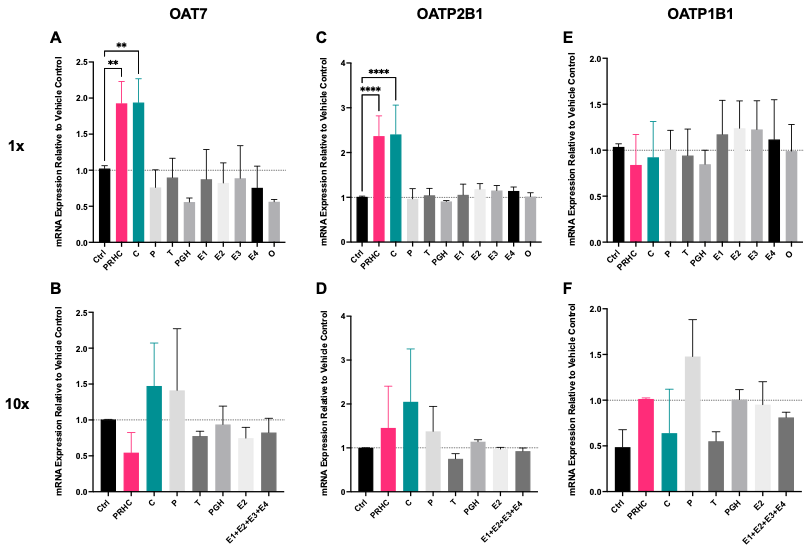

**Supplementary Figure 7. PRHC- and cortisol-mediated changes in OAT7, OATP2B1, and OATP1B1 mRNA expression in HepaRG cells.** HepaRG cells were treated for 72 h with vehicle, a pregnancy-related hormone cocktail (PRHC), or individual hormones at 1x (A, C, E) or 10x (B, D, F) their geometric mean third-trimester (T3) in vivo plasma concentrations. OAT7 (A, B), OATP2B1 (C, D), and OATP1B1 (E, F) mRNA expression, normalized to *GAPDH*, are expressed relative to vehicle (dotted line, 1.0). Where indicated, pooled estrogens are denoted E1+E2+E3+E4. At 1x T3 concentrations, PRHC and cortisol significantly increased OAT7 and OATP2B1 mRNA expression to a similar extent, but not OATP1B1 mRNA expression. At 10x T3 concentrations, none of the treatments significantly altered OAT7, OATP2B1, or OATP1B1 mRNA expression. Data are presented as mean ± SD from 3–5 independent experiments, each analyzed in technical triplicate. Statistical significance was assessed by one-way ANOVA followed by Dunnett’s multiple-comparison test versus vehicle (***P* < 0.01, *****P* < 0.0001). Ctrl, vehicle control; C, cortisol; P, progesterone; T, testosterone; PGH, placental growth hormone; E1, estrone; E2, estradiol; E3, estriol; E4, estetrol; O, oxytocin.

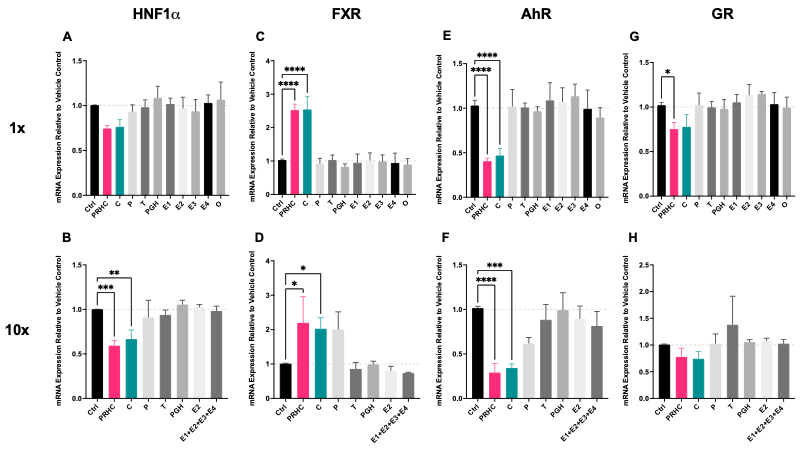

**Supplementary Figure 8. PRHC- and cortisol-mediated changes in HNF1α, FXR, AhR, and GR mRNA expression in HepaRG cells.** HepaRG cells were treated for 72 h with vehicle, a pregnancy-related hormone cocktail (PRHC), or individual hormones at 1x (A, C, E, G) or 10x (B, D, F, H) their geometric mean third-trimester (T3) *in vivo* plasma concentrations. HNF1α (A, B), FXR (C, D), AhR (E, F), and GR (G, H) mRNA expression, normalized to *GAPDH*, are expressed relative to vehicle control (dotted line, 1.0). Where indicated, pooled estrogens are denoted E1+E2+E3+E4. At 1x T3 concentrations, PRHC and cortisol significantly increased FXR mRNA expression and decreased AhR mRNA expression to a similar extent, whereas GR mRNA expression was modestly decreased by PRHC and HNF1α mRNA expression was not significantly altered. At 10x T3 concentrations, PRHC and cortisol decreased HNF1α and AhR mRNA expression and increased FXR mRNA expression, whereas GR mRNA expression was not significantly altered. Data are presented as mean ± SD from 3–5 independent experiments, each analyzed in technical triplicate. Statistical significance was assessed by one-way ANOVA followed by Dunnett’s multiple-comparison test versus vehicle (**P* < 0.05, ***P* < 0.01, ****P* < 0.001, *****P* < 0.0001). Ctrl, vehicle control; C, cortisol; P, progesterone; T, testosterone; PGH, placental growth hormone; E1, estrone; E2, estradiol; E3, estriol; E4, estetrol; O, oxytocin.

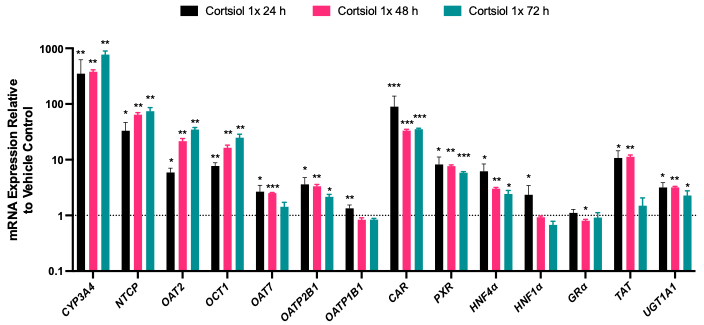

**Supplementary Figure 9. Time course of cortisol-mediated changes in hepatic transporter and regulatory factor mRNA expression in HepaRG cells.** HepaRG cells were treated with 1x cortisol for 24, 48, or 72 h. CYP3A4, NTCP, OAT2, OCT1, OAT7, OATP2B1, OATP1B1, CAR, PXR, HNF4α, HNF1α, GR, TAT, and UGT1A1 mRNA expression, normalized to *GAPDH*, are expressed relative to vehicle control (dotted line, 1.0). The 1x concentration corresponds to the geometric mean third-trimester (T3) *in vivo* plasma concentration of cortisol. Cortisol-responsive transporters and regulatory factors were altered to a similar extent by 48 and 72 h cortisol treatment, supporting the use of the 48 hr time point in subsequent mechanistic knockdown studies. Data are presented as mean ± SD of three replicates. Statistical significance was determined per mRNA target by one-way ANOVA followed by Dunnett’s multiple-comparison test versus vehicle (*P* < 0.05, **P* < 0.01, ***P* < 0.001).

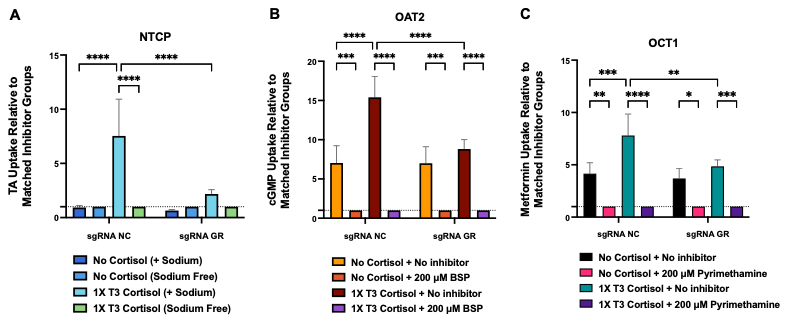

**Supplementary Figure 10. Cortisol increased NTCP-, OAT2-, and OCT1-mediated substrate uptake in sgRNA-NC-transfected HepaRG cells, and these increases were attenuated after GR knockdown.** HepaRG cells were transfected with sgRNA-NC or sgRNA-*NR3C1* and then treated with 1x T3 cortisol or vehicle for 48 h. Panel A shows taurocholic acid (TA) uptake by NTCP under sodium-containing or sodium-free conditions. Panel B shows cGMP uptake by OAT2 in the absence or presence of 200 μM BSP. Panel C shows metformin uptake by OCT1 in the absence or presence of 200 μM pyrimethamine. Uptake was measured using 50 nM [^3^H]TA, 40 nM [^3^H]cGMP, or 8.8 μM [^14^C]metformin, respectively, and is expressed relative to the matched inhibitor condition within each treatment group (dotted line, 1.0). Thus, uptake measured in the absence of inhibitor under vehicle conditions was normalized to the corresponding vehicle + inhibitor condition, and uptake measured in the absence of inhibitor under 1x T3 cortisol conditions was normalized to the corresponding 1x T3 cortisol + inhibitor condition. In sgRNA-NC-transfected cells, cortisol increased substrate uptake under non-inhibitory conditions for NTCP, OAT2, and OCT1. These cortisol-mediated increases were attenuated in sgRNA-*NR3C1*-transfected cells. Data are presented as mean ± SD of *n* = 3 independent experiments, each conducted in triplicate. Statistical significance was assessed by one-way ANOVA followed by Tukey’s multiple-comparison test (*P* < 0.05, **P* < 0.01, ***P* < 0.001, ****P* < 0.0001).

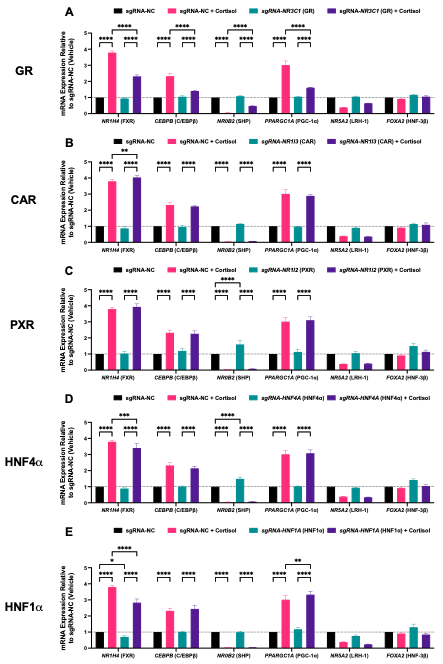

**Supplementary Figure 11. Exploratory analyses of additional cortisol-responsive regulatory factors after GR, CAR, PXR, HNF4α, or HNF1α knockdown in HepaRG cells.** HepaRG cells were transfected with sgRNA-NC or sgRNA-*NR3C1* (A), sgRNA-*NR1I3 (CAR)* (B), sgRNA-*NR1I2 (PXR)* (C), sgRNA-*HNF4A* (D), or sgRNA-*HNF1A* (E), and then treated with 1x T3 cortisol or vehicle for 48 h. *NR1H4* (FXR), *CEBPB* (C/EBPβ), *NR0B2* (SHP), *PPARGC1A* (PGC-1α), *NR5A2* (LRH-1), and *FOXA2* (HNF3β) mRNA expression, normalized to *GAPDH*, are expressed relative to sgRNA-NC + vehicle (dotted line, 1.0). Bars represent mean ± SD of *n* = 3 independent experiments, each conducted in triplicate. In sgRNA-NC-transfected cells, cortisol increased FXR, C/EBPβ, and PGC-1α mRNA expression and decreased SHP and LRH-1 mRNA expression, whereas HNF3β mRNA expression was minimally affected. GR knockdown attenuated these cortisol-mediated changes. CAR or PXR knockdown largely preserved this cortisol-responsive profile, although CAR knockdown modestly enhanced FXR induction. HNF4α knockdown modestly attenuated FXR induction, whereas HNF1α knockdown attenuated FXR induction and modestly enhanced PGC-1α induction. Statistical significance was assessed by one-way ANOVA followed by Tukey’s multiple-comparison test (**P* < 0.05, ***P* < 0.01, ****P* < 0.001, *****P* < 0.0001).

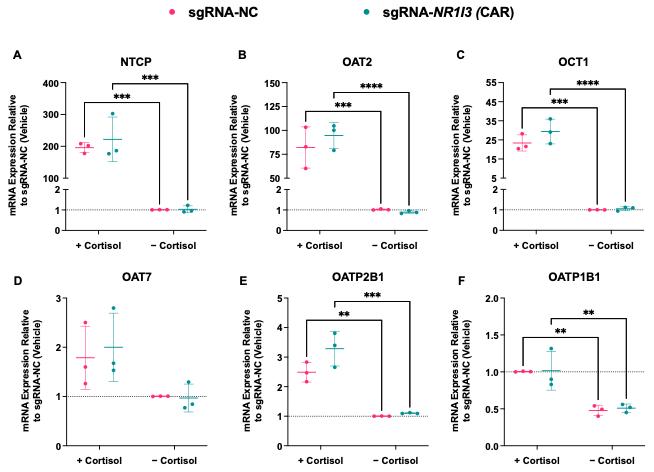

**Supplementary Figure 12. CAR knockdown does not significantly alter cortisol-mediated transporter mRNA expression in HepaRG cells.** HepaRG cells were transfected with sgRNA-NC or sgRNA-*NR1I3* and then treated with 1x T3 cortisol or vehicle for 48 h. NTCP (A), OAT2 (B), OCT1 (C), OAT7 (D), OATP2B1 (E), and OATP1B1 (F) mRNA expression, normalized to *GAPDH*, are expressed relative to sgRNA-NC + vehicle (dotted line, 1.0). Points denote independent experiments; horizontal lines and error bars represent mean ± SD of *n* = 3 independent experiments, each conducted in triplicate. In sgRNA-NC-transfected cells, cortisol increased NTCP, OAT2, OCT1, and OATP2B1 mRNA expression and decreased OATP1B1 mRNA expression. These cortisol-mediated changes were preserved after CAR knockdown. Transporter mRNA expression in cortisol-treated sgRNA-*NR1I3* cells was not significantly different from that in cortisol-treated sgRNA-NC cells. Statistical significance was assessed by one-way ANOVA followed by Tukey’s multiple-comparison test (***P* < 0.01, ****P* < 0.001, *****P* < 0.0001).

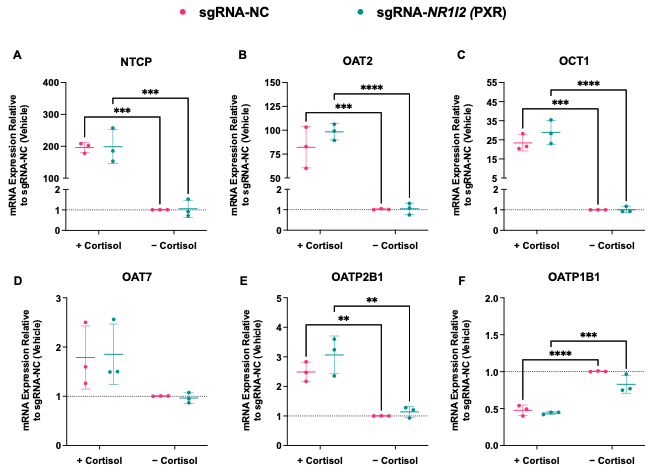

**Supplementary Figure 13. PXR knockdown does not significantly alter cortisol-mediated transporter mRNA expression in HepaRG cells.** HepaRG cells were transfected with sgRNA-NC or sgRNA-*NR1I2* and then treated with 1x T3 cortisol or vehicle for 48 h. NTCP (A), OAT2 (B), OCT1 (C), OAT7 (D), OATP2B1 (E), and OATP1B1 (F) mRNA expression, normalized to *GAPDH*, are expressed relative to sgRNA-NC + vehicle (dotted line, 1.0). Points denote independent experiments; horizontal lines and error bars represent mean ± SD of *n* = 3 independent experiments, each conducted in technical triplicate. In sgRNA-NC-transfected cells, cortisol increased NTCP, OAT2, OCT1, and OATP2B1 mRNA expression and decreased OATP1B1 mRNA expression. These cortisol-mediated changes were preserved after PXR knockdown. Transporter mRNA expression in cortisol-treated sgRNA-*NR1I2* cells was not significantly different from that in cortisol-treated sgRNA-NC cells. Statistical significance was assessed by two-way ANOVA followed by Tukey’s multiple-comparison test (**P* < 0.01, ***P* < 0.001, ****P* < 0.0001).

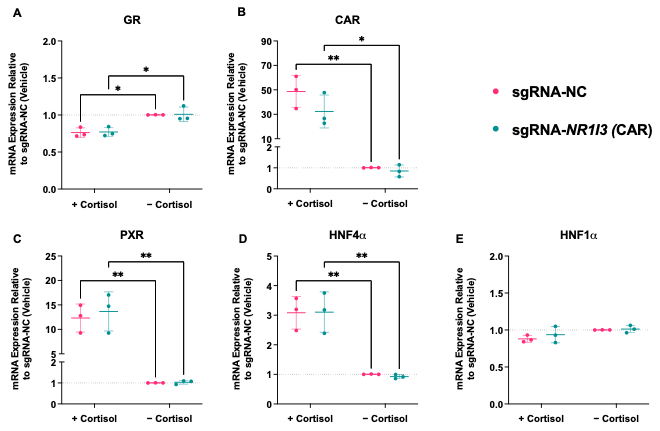

**Supplementary Figure 14. CAR knockdown does not significantly alter cortisol-mediated regulatory factor mRNA expression in HepaRG cells.** HepaRG cells were transfected with sgRNA-NC or CAR-targeting sgRNA (sgRNA-*NR1I3*) and then treated with 1x T3 cortisol or vehicle for 48 h. GR (A), CAR (B), PXR (C), HNF4α (D), and HNF1α (E) mRNA expression, normalized to *GAPDH*, are expressed relative to sgRNA-NC + vehicle (dotted line, 1.0). Points denote independent experiments; horizontal lines and error bars represent mean ± SD of *n* = 3 independent experiments, each conducted in technical triplicate. In sgRNA-NC-transfected cells, cortisol decreased GR mRNA expression and increased CAR, PXR, and HNF4α mRNA expression, whereas HNF1α mRNA expression was not significantly altered. These cortisol-mediated changes were preserved after CAR knockdown, and regulatory factor mRNA expression in cortisol-treated sgRNA-*NR1I3* cells was not significantly different from that in cortisol-treated sgRNA-NC cells. Statistical significance was assessed by two-way ANOVA followed by Tukey’s multiple-comparison test (**P* < 0.05, ***P* < 0.01).

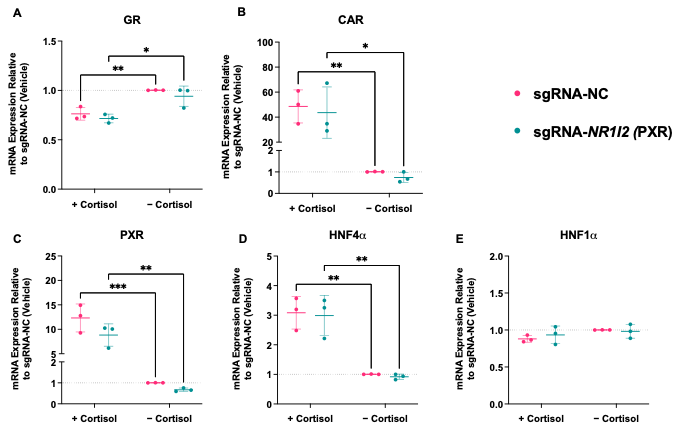

**Supplementary Figure 15. PXR knockdown does not significantly alter cortisol-mediated regulatory factor mRNA expression in HepaRG cells.** HepaRG cells were transfected with sgRNA-NC or PXR-targeting sgRNA (sgRNA-*NR1I2*) and then treated with 1x T3 cortisol or vehicle for 48 h. GR (A), CAR (B), PXR (C), HNF4α (D), and HNF1α (E) mRNA expression, normalized to *GAPDH*, are expressed relative to sgRNA-NC + vehicle (dotted line, 1.0). Points denote independent experiments; horizontal lines and error bars represent mean ± SD of *n* = 3 independent experiments, each conducted in triplicate. In sgRNA-NC-transfected cells, cortisol decreased GR mRNA expression and increased CAR, PXR, and HNF4α mRNA expression, whereas HNF1α mRNA expression was not significantly altered. These cortisol-mediated changes were preserved after PXR knockdown, and mRNA expression of these regulatory factors in cortisol-treated sgRNA-*NR1I2* cells was not significantly different from that in cortisol-treated sgRNA-NC cells. Statistical significance was assessed by two-way ANOVA followed by Tukey’s multiple-comparison test (**P* < 0.05, ***P* < 0.01).

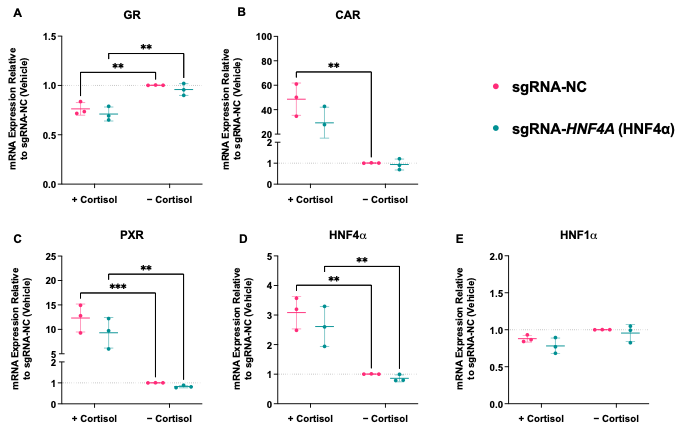

**Supplementary Figure 16. HNF4α knockdown does not significantly alter cortisol-mediated regulatory factor mRNA expression in HepaRG cells.** HepaRG cells were transfected with sgRNA-NC or HNF4α-targeting sgRNA (sgRNA-*HNF4A*) and then treated with 1x T3 cortisol or vehicle for 48 h. GR (A), CAR (B), PXR (C), HNF4α (D), and HNF1α (E) mRNA expression, normalized to *GAPDH*, are expressed relative to sgRNA-NC + vehicle (dotted line, 1.0). Points denote independent experiments; horizontal lines and error bars represent mean ± SD of *n* = 3 independent experiments, each conducted in technical triplicate. In sgRNA-NC-transfected cells, cortisol decreased GR mRNA expression and increased CAR, PXR, and HNF4α mRNA expression, whereas HNF1α mRNA expression was not significantly altered. These cortisol-mediated changes were preserved after HNF4α knockdown, and regulatory factor mRNA expression in cortisol-treated sgRNA-*HNF4A* cells was not significantly different from that in cortisol-treated sgRNA-NC cells. Statistical significance was assessed by two-way ANOVA followed by Tukey’s multiple-comparison test (***P* < 0.01, ****P* < 0.001).

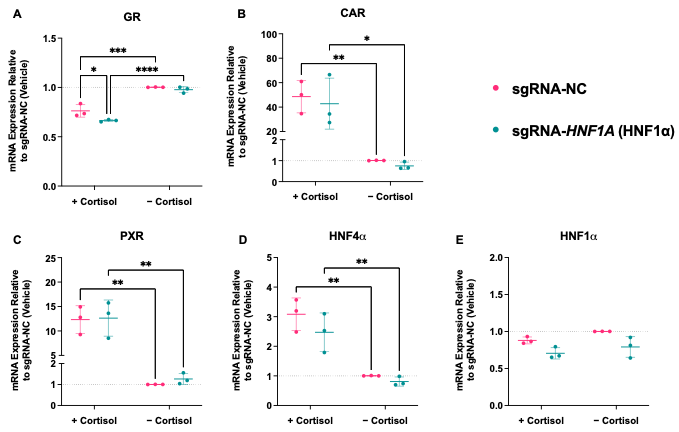

**Supplementary Figure 17. HNF1α knockdown further decreases GR mRNA expression but does not significantly alter other cortisol-mediated regulatory factor mRNA changes in HepaRG cells.** HepaRG cells were transfected with sgRNA-NC or HNF1α-targeting sgRNA (sgRNA-*HNF1A*) and then treated with 1x T3 cortisol or vehicle for 48 h. GR (A), CAR (B), PXR (C), HNF4α (D), and HNF1α (E) mRNA expression, normalized to *GAPDH*, are expressed relative to sgRNA-NC + vehicle (dotted line, 1.0). Points denote independent experiments; horizontal lines and error bars represent mean ± SD of *n* = 3 independent experiments, each conducted in triplicate. In sgRNA-NC-transfected cells, cortisol decreased GR mRNA expression and increased CAR, PXR, and HNF4α mRNA expression, whereas HNF1α mRNA expression was not significantly altered. HNF1α knockdown further decreased GR mRNA expression in cortisol-treated cells, but did not significantly alter cortisol-mediated changes in CAR, PXR, HNF4α, or HNF1α mRNA expression. Statistical significance was assessed by two-way ANOVA followed by Tukey’s multiple-comparison test (**P* < 0.05, ***P* < 0.01, ****P* < 0.001, *****P* < 0.0001).

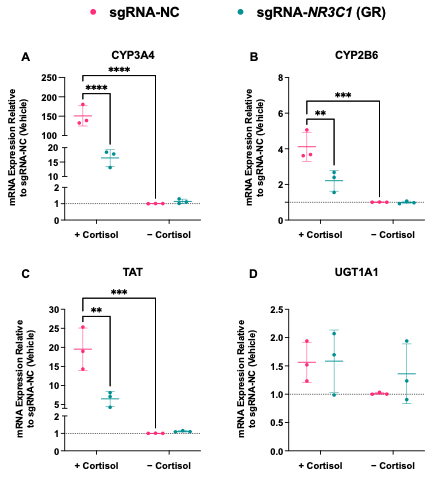

**Supplementary Figure 18. GR knockdown selectively attenuates cortisol-mediated induction of CYP3A4, CYP2B6, and TAT mRNA in HepaRG cells.** HepaRG cells were transfected with sgRNA-NC or *NR3C1*-targeting sgRNA (sgRNA-*NR3C1*), then treated with cortisol or vehicle for 48 h. mRNA expression of CYP3A4 (A), CYP2B6 (B), TAT (C), and UGT1A1 (D) was quantified by RT-qPCR, normalized to *GAPDH*, and expressed relative to sgRNA-NC vehicle control. Cortisol induced CYP3A4 (151-fold), CYP2B6 (4-fold), TAT (20-fold), and UGT1A1 (1.6-fold) mRNA expression in sgRNA-NC cells, and GR knockdown markedly attenuated these responses. Data are shown as mean ± SD from three independent experiments. Statistical significance was assessed by two-way ANOVA with multiple-comparison correction (**P<0.01, ***P<0.001, ****P<0.0001).

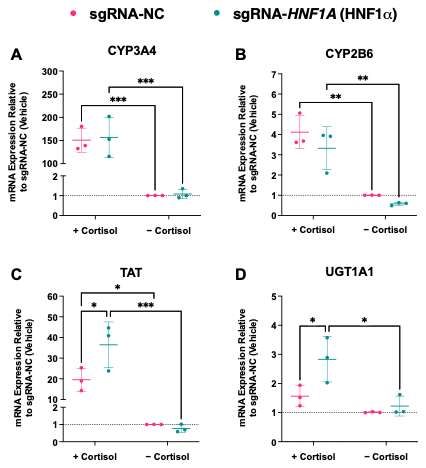

**Supplementary Figure 19. HNF1α knockdown enhances cortisol-mediated induction of TAT and UGT1A1 mRNA in HepaRG cells.** HepaRG cells were transfected with sgRNA-NC or *HNF1A*-targeting sgRNA (sgRNA-*HNF1A*), then treated with cortisol or vehicle for 48 h. mRNA expression of CYP3A4 (A), CYP2B6 (B), TAT (C), and UGT1A1 (D) was quantified by RT-qPCR, normalized to *GAPDH*, and expressed relative to sgRNA-NC vehicle control. HNF1α knockdown enhanced cortisol-induced TAT (+86%) and UGT1A1 (+81%) mRNA expression but did not blunt CYP3A4 and CYP2B6 induction. Data are shown as mean ± SD from three independent experiments. Statistical significance was assessed by two-way ANOVA with multiple-comparison correction (*P<0.05, **P<0.01, ***P<0.001).

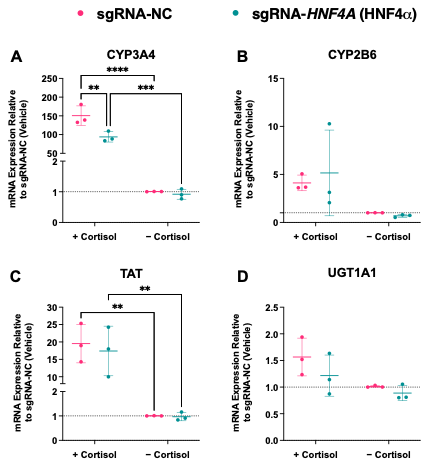

**Supplementary Figure 20. HNF4α knockdown selectively attenuates cortisol-mediated induction of CYP3A4 mRNA in HepaRG cells.** HepaRG cells were transfected with sgRNA-NC or *HNF4A*-targeting sgRNA (sgRNA-*HNF4A*), then treated with cortisol or vehicle for 48 h. mRNA expression of CYP3A4 (A), CYP2B6 (B), TAT (C), and UGT1A1 (D) was quantified by RT-qPCR, normalized to *GAPDH*, and expressed relative to sgRNA-NC vehicle control. HNF4α knockdown selectively reduced cortisol-induced CYP3A4 mRNA expression (−38%). Data are shown as mean ± SD from three independent experiments. Statistical significance was assessed by two-way ANOVA with multiple-comparison correction (**P<0.01, ***P<0.001, ****P<0.0001).

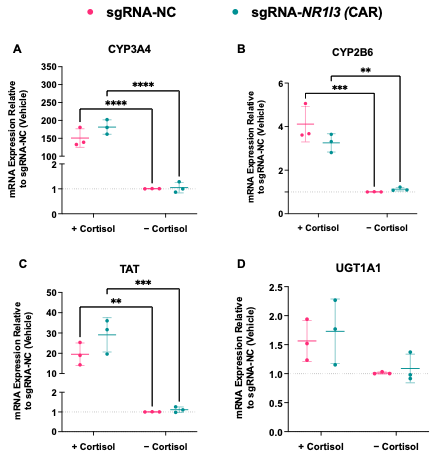

**Supplementary Figure 21. CAR knockdown did not significantly alter cortisol-mediated induction of CYP3A4, CYP2B6, and TAT mRNA in HepaRG cells.** HepaRG cells were transfected with sgRNA-NC or *NR1I3*-targeting sgRNA (sgRNA-*NR1I3*), then treated with cortisol or vehicle for 48 h. mRNA expression of CYP3A4 (A), CYP2B6 (B), TAT (C), and UGT1A1 (D) was quantified by RT-qPCR, normalized to *GAPDH*, and expressed relative to sgRNA-NC vehicle control. CAR knockdown did not significantly alter the cortisol-mediated induction of these control genes. Data are shown as mean ± SD from three independent experiments. Statistical significance was assessed by two-way ANOVA with multiple-comparison correction (**P<0.01, ***P<0.001, ****P<0.0001).

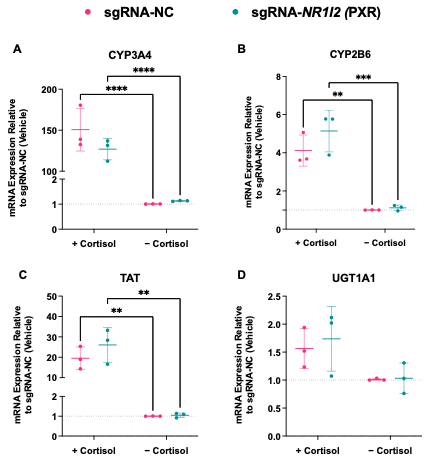

**Supplementary Figure 22. PXR knockdown did not significantly alter cortisol-mediated induction of CYP3A4, CYP2B6, and TAT mRNA in HepaRG cells.** HepaRG cells were transfected with sgRNA-NC or *NR1I2*-targeting sgRNA (sgRNA-*NR1I2*), then treated with cortisol or vehicle for 48 h. mRNA expression of CYP3A4 (A), CYP2B6 (B), TAT (C), and UGT1A1 (D) was quantified by RT-qPCR, normalized to *GAPDH*, and expressed relative to sgRNA-NC vehicle control. PXR knockdown did not significantly alter the cortisol-mediated induction of these control genes. Data are shown as mean ± SD from three independent experiments. Statistical significance was assessed by two-way ANOVA with multiple-comparison correction (**P<0.01, ***P<0.001, ****P<0.0001).

**Supplementary Table 1. sgRNA sequences used for CRISPR-Cas9-mediated knockdown in HepaRG cells.**

| **Target gene** | **Encoded protein** | **ThermoFisher TrueGuide ID** | **sgRNA sequence, 5′**–**3′** |
| --- | --- | --- | --- |
| ***NR3C1*** | GR | CRISPR633788_SGM | GCCAAATCAGCCTTTCCTCG |
| ***HNF1A*** | HNF1α | CRISPR947755_SGM | GATGTTGTGCTGCTGCAGGT |
| ***HNF4A*** | HNF4α | CRISPR732495_SGM | GTCCATGTCGACGAGGGTTT |
| ***NR1I2*** | PXR | CRISPR821485_SGM | GTCAACGCAGATGAGGAAGT |
| ***NR1I3*** | CAR | CRISPR743801_SGM | CGCATTAAAGTGGTAGCCTG |

**Supplementary Table 2. PCR primers used to amplify genomic regions flanking CRISPR-Cas9 target sites for cleavage detection**

| **Target gene and encoded protein** | **Primer Sequence** | **Expected cleavage fragments (bp)** |
| --- | --- | --- |
| *NR3C1* (GR) | Forward (5′–3′): ACCCTCACTGGCTGTCGCTTCT  Reverse (5′–3′): ATTGCCACCGTTGGTGCCAGTC | 205/201 |
| *HNF1A* (HNF1α) | Forward (5′–3′): TCATGCACAGTCCCCACCCTCA  Reverse (5′–3′): ATGGTGAAGCTTCCAGCCCCCA | 155/245 |
| *HNF4A* (HNF4α) | Forward (5′–3′): AACAGCCAAATCCCTGCAGCCC  Reverse (5′–3′): ACTGGCACACCTGGGCACATCT | 274/133 |
| *NR1I3* (CAR) | Forward (5′–3′): TGGGCAACATGGCAAAACCCCA  Reverse (5′–3′): TGTAGCTGGACAGGCTTGGGCA | 283/127 |
| *NR1I2* (PXR) | Forward (5′–3′): ATTCCAACCCCCATTCCCCGCT  Reverse (5′–3′): ACGTGGGTGAAGGCTGATGGGT | 431/160 |
